# A Flexible Model of Working Memory

**DOI:** 10.1101/407700

**Authors:** Flora Bouchacourt, Timothy J. Buschman

## Abstract

Working memory is fundamental to cognition, allowing one to hold information ‘in mind’ and use it to guide behavior. A defining characteristic of working memory is its flexibility: we can hold anything in mind. However, typical models of working memory rely on finely tuned, content-specific, attractors to persistently maintain neural activity and therefore do not allow for the flexibility observed in behavior. Here we present a flexible model of working memory that maintains representations through random recurrent connections between two layers of neurons: a structured ‘sensory’ layer and a randomly connected, unstructured, layer. As the interactions are untuned with respect to the content being stored, the network is able to maintain any arbitrary input. However, this flexibility comes at a cost: the random connections overlap, leading to interference between representations and limiting the memory capacity of the network. Additionally, our model captures several other key behavioral and neurophysiological characteristics of working memory.

## Introduction

Working memory is the ability to hold information ‘in mind’. It acts as a workspace on which information can be held, manipulated, and then used to guide behavior. In this way, working memory plays a critical role in cognition, allowing behavior to be decoupled from the immediate sensory world. However, despite its importance, the circuit mechanisms that support working memory remain unclear. In particular, existing models fail to capture key behavioral and neural characteristics of working memory.

Working memory has two defining behavioral characteristics. First, it is highly flexible: one can hold anything in mind and can do it from the first experience. This flexibility allows working memory to act as a generic workspace to support cognition. Second, working memory has a severely limited capacity. Humans and monkeys are able to maintain only 3-4 objects at once [Buschman et al., 2011; Cowan, 2010; Luck and Vogel, 1997; Ma et al., 2014]. In other words, although one can hold anything in mind, one can only hold a few of them at a time.

In addition to these behavioral characteristics, previous work has identified several neural characteristics of working memory. First, the contents of working memory are thought to be represented in both the persistent activity of neurons [Funahashi et al., 1989; Fuster, 1999; Goldman-Rakic, 1995; Miller et al., 1993; Romo et al., 2002] and in the dynamic evolution of neural activity over time [Barak et al., 2010; Jun et al., 2010; Murray et al., 2016; Rainer et al., 1998; Stokes, 2015]. Second, working memory representations are distributed across the brain [Christophel et al., 2017]: working memory representations have been observed in prefrontal [Ester et al., 2015; Riley and Constantinidis, 2016], parietal [Christophel et al., 2015; Constantinidis and Steinmetz, 1996], and sensory cortex [Harrison and Tong, 2009; Pasternak and Greenlee, 2005; Serences et al., 2009], as well as sub-cortical regions such as the basal ganglia [Alexander, 1987] and thalamus [Watanabe and Funahashi, 2012]. Third, increasing the number of items held in working memory (the ‘memory load’), increases the overall activity in these brain regions (up to an individual capacity limit; [Vogel and Machizawa, 2004]). However, increasing memory load also reduces the selectivity of individual neurons in a divisive-normalization-like manner; the firing rate of neurons selective for one item is decreased with the addition of more items. This is thought to underlie the reduction in memory performance and accuracy at high memory loads [Buschman et al., 2011; Sprague et al., 2014].

Theoretical models have captured some, but not all, of these characteristics. The dominant model of working memory is that persistent neural activity is the result of recurrent network interactions, either within or between brain regions [Barak and Tsodyks, 2014; Brunel, 2003; Compte et al., 2000; Machens et al., 2005; Wang, 2001]. These models have a limited capacity, as lateral inhibition limits the number of simultaneous patterns of activity that can be maintained [Edin et al., 2009; Swan and Wyble, 2014]. However, these models are inflexible. They rely on fine tuning of connections to embed stable fixed points in the network dynamics specific to the content being stored. For example, heading position is represented by a ring attractor network [Kim et al., 2017; Seelig and Jayaraman, 2015]. These connections must either be hardwired or learned for each type of information, and so the network is not able to flexibly represent novel, unexpected, stimuli.

Models that represent working memory as a result of transient dynamics in neural activity have similar constraints on flexibility. They require learning to embed the dynamics, to decode the temporally evolving representations, or to fine tune connections such that the dynamics are orthogonal to mnemonic representations [Brody et al., 2003; Druckmann and Chklovskii, 2012; Jaeger, 2002; Rajan et al., 2016; Seung et al., 2000; Vogels et al., 2005].

Other models capture the flexibility of working memory, such as those that hypothesize working memory representations are encoded in short-term synaptic plasticity or changes in single-cell biophysics [Egorov et al., 2002; Fransén et al., 2006; Hasselmo and Stern, 2006; Loewenstein and Sompolinsky, 2003; Mongillo et al., 2008]. However, these models do not directly explain the limited capacity of working memory and do not capture many of the neurophysiological characteristics of working memory, such as the coexistence of persistent and dynamic representations seen in neural data.

Here we propose a flexible model of working memory that relies on random reciprocal connections to generate persistent activity. As the connections are random, they are inherently untuned with respect to the content being stored and do not need to be learned, allowing the network to maintain any representation. However, this flexibility comes at a cost - when multiple memories are stored in the network, they begin to interfere, resulting in divisive-normalization-like reduction of responses and imposing a capacity limit on the network. Thus, our model provides a mechanistic explanation for the limited capacity of working memory; it is a necessary trade-off for its flexibility.

## Results

### A flexible model of working memory

We model a simplified two-layer network of Poisson spiking neurons (Fig. 1A; see Methods for a detailed description). The first layer is the ‘sensory network’ and consists of 8 independent ring-like sub-networks (each modeled here with 512 neurons). These sub-networks mimic simplified sensory networks and can be thought of as encoding the identity of stimuli at different locations in space. Therefore, we can vary working memory load by varying the number of sensory sub-networks receiving inputs.

**Figure 1.**
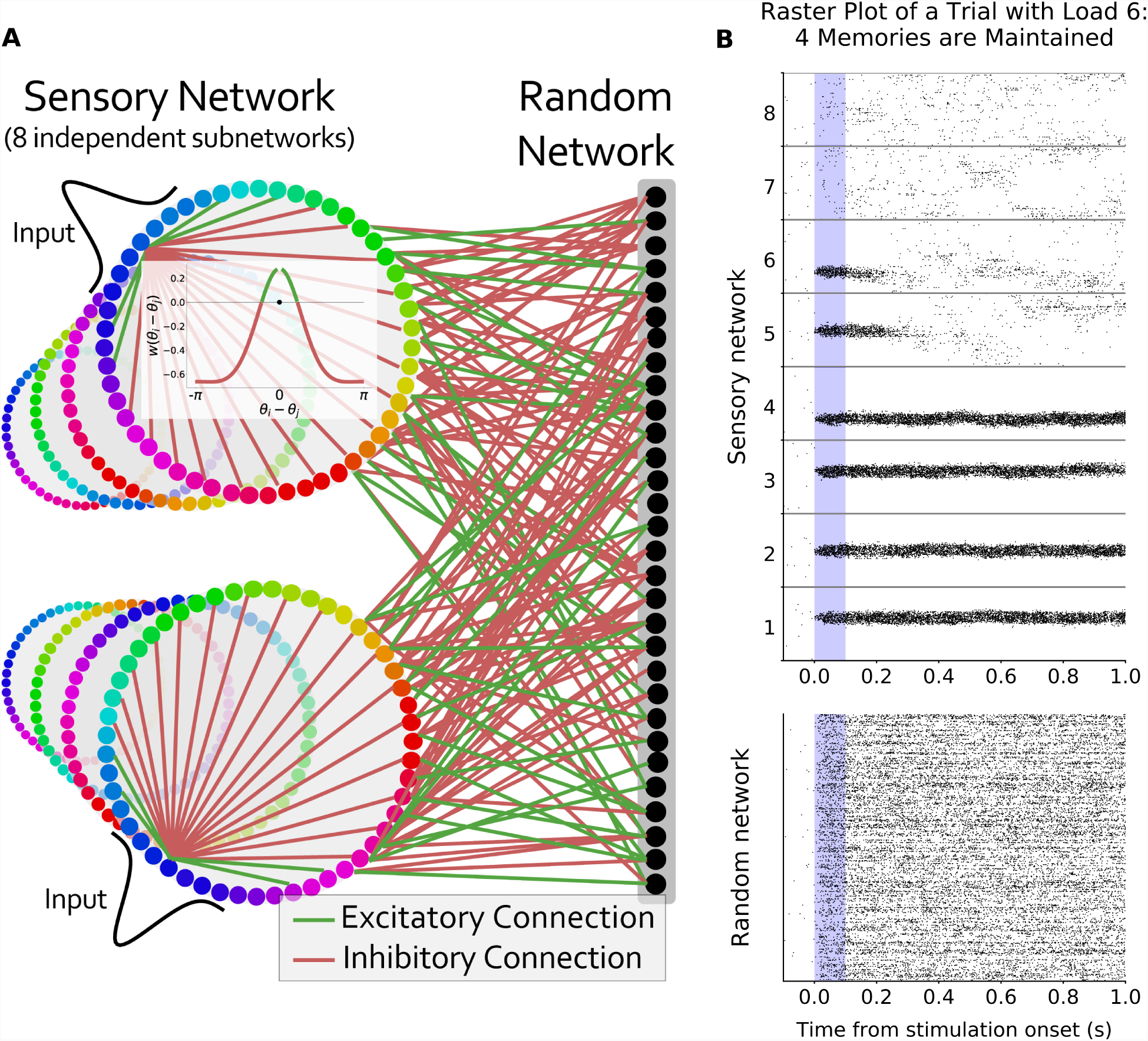
Flexible working memory through interactions between a structured network and a random network. **(A)** Model layout. The sensory network is composed of 8 ring-like sub-networks (although other architectures can be used, see Fig. S8). The inset represents the center-surround connectivity structure of a sensory sub-network (excitatory connections in green, inhibitory connections in red). The connections to the random network are randomly assigned and balanced. The unstructured nature of these connections endows the network with its flexibility. **(B)** Raster plot of an example trial with 8 sensory sub-networks (512 neurons each) randomly connected to the same random network (1024 neurons). Six sensory sub-networks receive a Gaussian input for 0.1 seconds during the ‘stimulus presentation’ period (shaded blue region). Representations are maintained (without external drive) for four of the inputs.

Neurons within each sensory sub-network are arranged topographically according to selectivity. Position around the ring corresponds to specific values of an encoded feature, such as orientation or color. Consistent with biological observations [Funahashi et al., 1989; Kiyonaga and Egner, 2016; Kuffler, 1953], connections within a sensory sub-network have a center-surround structure: neurons with similar selectivity share excitatory connections while inhibition is broader (Fig. 1A, inset). However, recurrent excitation within each sub-network is too low to maintain memories by themselves. For simplicity, each sensory sub-network is independent (i.e. there are no connections between them).

The second layer is the ‘random network’ (modeled here with 1024 neurons). Neurons in this layer are randomly and reciprocally connected to neurons in the sensory sub-networks. Each neuron in the random network has bi-directional excitatory connections with a random subset of neurons in the sensory network (with a likelihood *γ*; here, 0.35). Importantly, all sensory neurons converge onto the same random network. The connections between the sensory and random networks are balanced, such that individual neurons receive an equal amount of excitatory and inhibitory drive (i.e. the summed input weight to each neuron in the random network from the sensory network is zero; as are connections to the sensory network from the random network). To achieve this, all pairs of random and sensory neurons without excitatory connections have direct, weak, inhibitory connections, such that the sum of excitatory weights equals the sum of inhibitory weights for any given neuron (see Methods for details). Such excitation-inhibition balance is consistent with neurophysiological findings [Mariño et al., 2005; Vogels and Abbott, 2005; Vogels et al., 2011].

Despite this simple architecture, the network is able to maintain stimulus inputs over an extended memory delay (Fig. 1B). This is due to the bi-directional and reciprocal connections between the sensory and random networks. Activity in the sensory network feeds-forward into the random network, activating a random subset of neurons (Fig. 1B, bottom). In turn, neurons from the random network feedback into the sensory network, maintaining activity after the stimulus input is removed (Fig. 1B, top, sensory sub-networks 1-4). The reciprocal nature of the connections ensures the synaptic drive fed back into a sensory sub-network from the random network closely matches its own representation (Fig. S1). In this way, the network can flexibly maintain the representation of any input into the sensory network.

Neurons from the random network project back to multiple sensory sub-networks. However, activity in the random network does not lead to sustained representations in other sensory sub-networks. As feedback connections are random and balanced they ‘destructively interfere’ at other locations (i.e. the expected value of feedback projections is zero). Therefore, there is a slight increase in noise but no spuriously sustained representations (Fig. 1B, top, sensory sub-networks 7 and 8). In other words, the feedback input from the random network to other sensory sub-networks is orthogonal to their inherent dynamics and therefore is not sustained.

Neurons in the sensory network show physiologically realistic tuning curves, due to their center-surround architecture (as illustrated in Fig. 2A,B). This tuning is effectively inherited by the random network, although with greater complexity (Fig. 2C,D), matching neurophysiological findings [Funahashi et al., 1989; Mendoza-Halliday et al., 2014; Zaksas and Pasternak, 2006]. However, there is no consistency in the tuning of neurons in the random network across inputs to different sensory networks (Fig. 2E). This is due to the fact that connectivity is random and therefore inconsistent across sensory networks. This leads to neurons in the random network showing ‘conjunctive’ coding (Fig. 2F), preferentially responding to different input values for different sensory sub-networks (e.g. different colors at different locations, as is observed in prefrontal neurons) and responding to unique combinations of inputs into multiple sensory sub-networks [Fusi et al., 2016; Rigotti et al., 2013].

**Figure 2.**
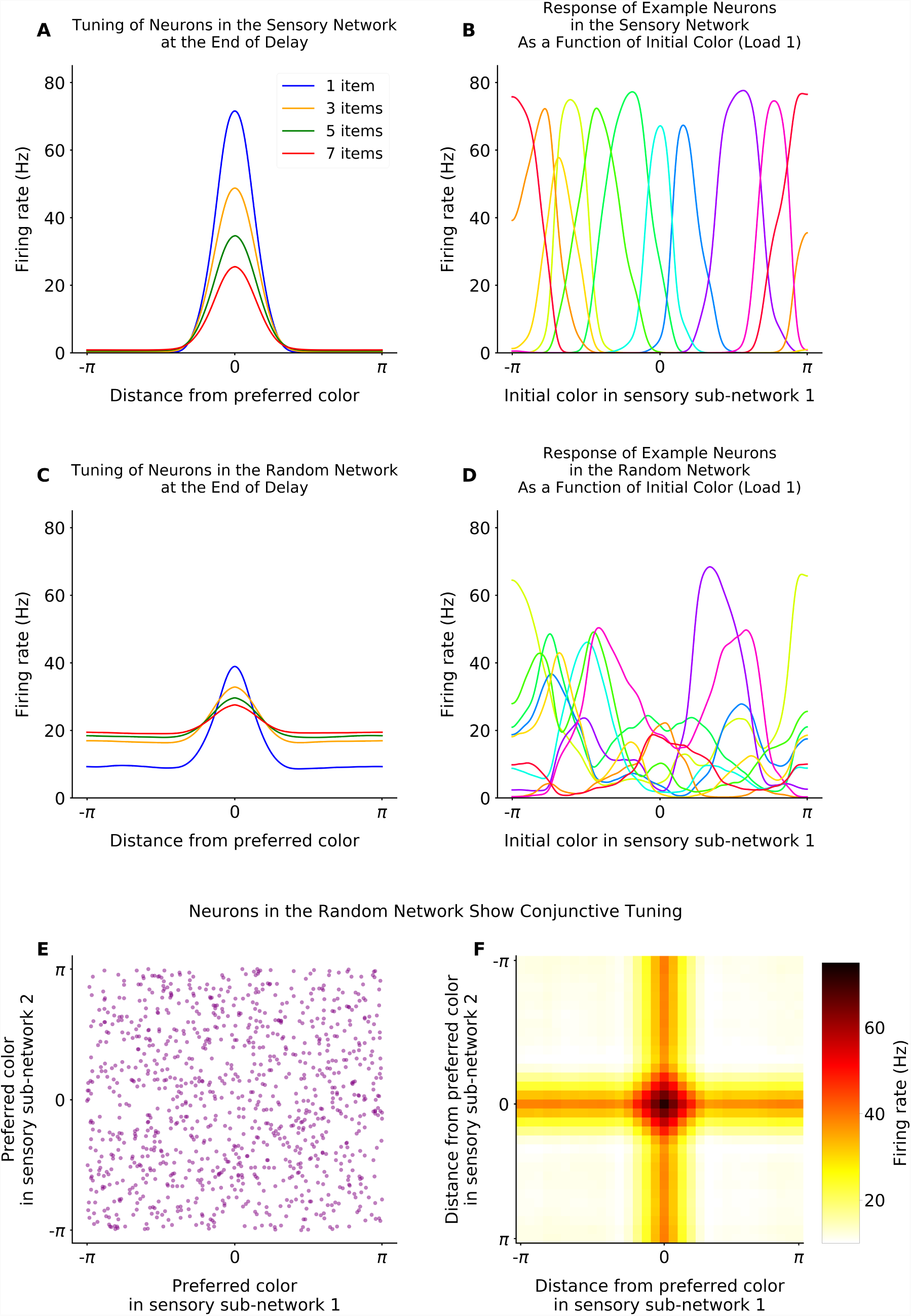
Tuning in the sensory and random networks. **(A)** Neurons in the sensory sub-networks have physiologically realistic tuning curves. Response of neurons (y-axis) is shown at the end of the delay period, relative to each neuron’s preferred stimulus input (x-axis). Tuning decreases with increased working memory load (colored lines). **(B)** Example tuning curves of a subset of neurons in the sensory network, distributed across space. **(C)** Neurons in the random network show physiologically realistic tuning, inherited from the sensory network. As in **A**, tuning is shown relative to the preferred stimulus input. As in the sensory network, tuning decreases with memory load (colored lines). **(D)** As in **B**, example tuning curves of a subset of neurons in the random network. **(E and F)** Neurons in the random network show conjunctive tuning. First, as seen in **E**, neurons respond to the conjunction of stimulus identity and location. The preferred stimulus of neurons from the random network is uncorrelated between sensory sub-networks, due to the random projections between networks (shown here as preferred input for sensory sub-network 1, x-axis, and sub-network 2, y-axis). Second, as seen in **F**, neurons in the random network preferentially respond to a conjunction of stimulus inputs across sensory networks. This is shown here by the two-dimensional tuning curve of neurons from the random network when sensory sub-network 1 and sensory sub-network 2 receive an input during stimulation. The firing rate (color axis) is aligned on the x-axis with respect to the preferred input for sensory sub-network 1, and on the y-axis with respect to the preferred input for sensory sub-network 2, showing a peaked response at the conjunction of the two stimuli.

### Interference between memory representations imposes a capacity limit

Multiple memories can be stored in the network simultaneously (Fig. 1B). For a few items (typically < 3), memories do not significantly interfere - there is sufficient space in the high-dimensional random network for multiple patterns to be maintained. However, as the number of inputs to the sensory networks is increased, interference in the random network increases, eventually causing memory failures (seen in Fig. 1B, sensory sub-networks 5 and 6). Figure 3A shows the percentage of correct memories at the end of the delay period, as a function of load (e.g. number of items presented; see Methods for details). This closely matches behavioral results in both humans and monkeys [Buschman et al., 2011; Cowan, 2010; Luck and Vogel, 1997]. Also consistent with behavior, the speed of forgetting during the delay period increases with load (Fig. 3B). As with all of our results, this is not due to tuning of network parameters; parameters were set to maximize the total number of items that were remembered across all loads, not to match any behavioral or electrophysiological results (see Methods).

**Figure 3.**
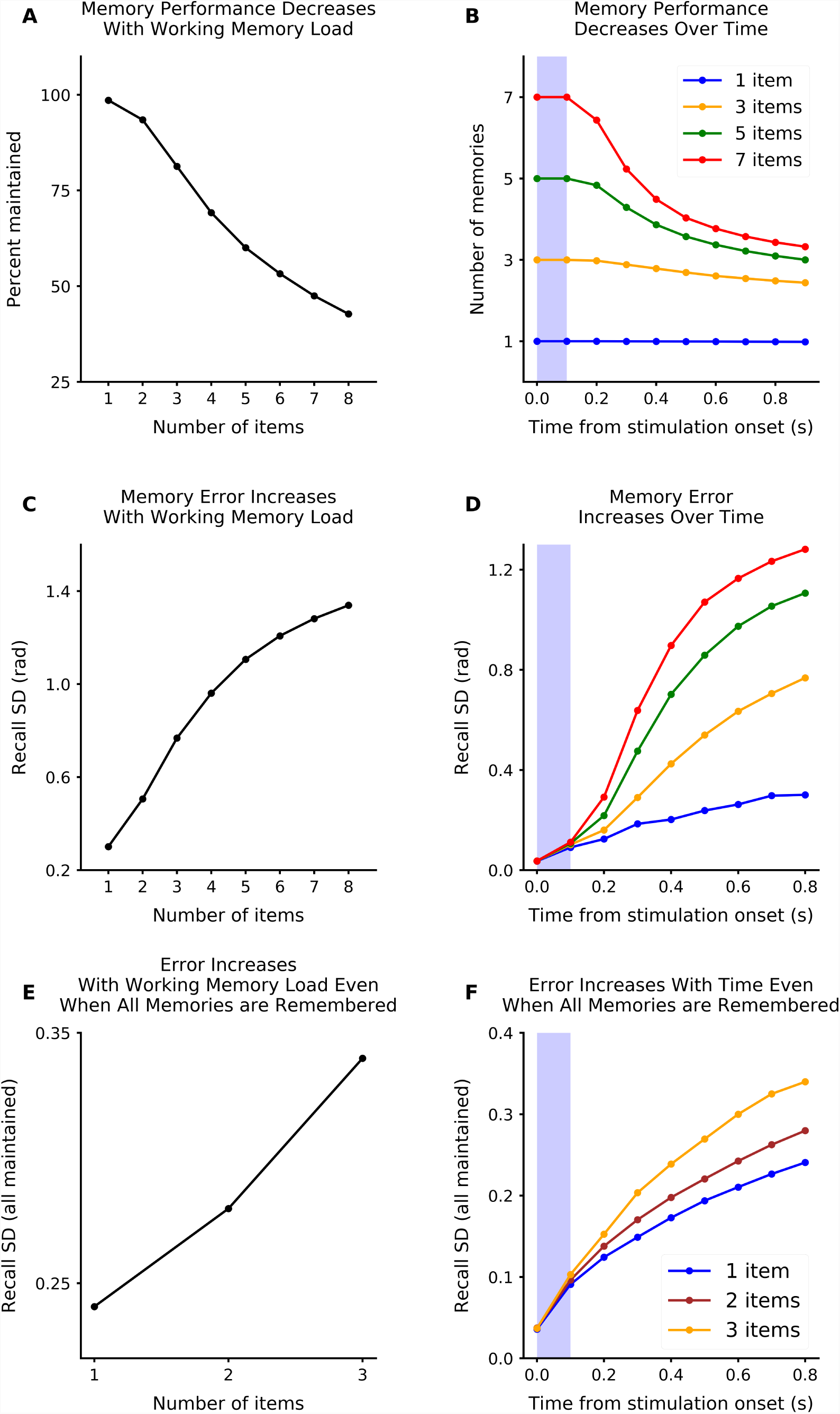
Memory performance decreases with working memory load. **(A)** The network has a limited memory capacity. The percentage of memories maintained (measured as memory representations *≥* 3 Hz after the delay period, see Methods for details) decreases with memory load (‘number of items’). **(B)** The speed of forgetting during the delay period is load-dependent. There is a sharper decrease in memory performance for higher memory loads. **(C)** Memory precision decreases with working memory load. The precision was measured as the standard deviation of the circular error computed from the maximum likelihood estimate of the memory from the sensory network (decoding time window lasts 0.1s and started 0.8s after the stimulus onset). **(D)** Memory precision decreases over time and with load. Decoding time window is 0.1s forward from the time point referenced. **(E-F)** Memory precision decreases with memory load, even when all memories are maintained. This shows that the decay in precision is not entirely due to forgetting. The panels display the same measure as in **C-D**, respectively, but only the simulations where all memories are maintained are considered.

Consistent with behavioral studies [Bays et al., 2009; Ma et al., 2014; Pertzov et al., 2013, 2017; Rademaker et al., 2018; Zhang and Luck, 2008], errors in working memory increased with the number of items held in memory (Fig. 3C and S2A). As with forgetting, these errors increased over time in a load-dependent manner (Fig. 3D and S2B). The increase in error was not just due to forgetting of inputs – even when an input was successfully maintained, there was an increase in error with increasing memory load (Fig. 3E,F). This is consistent with behavioral studies that have shown increases in memory error, even below working memory capacity limits [Bays et al., 2009; Ma et al., 2014; Pertzov et al., 2017; Rademaker et al., 2018].

Our model provides a simple, mechanistic explanation for the limited capacity of working memory: it is due to interference in neural representations in the shared random network. This is a natural consequence of the convergent, random connectivity between the two non-linear networks; as multiple inputs are presented to the sensory networks, their representations interfere in the random network, disrupting maintenance of representations. This is an unavoidable consequence of the convergence and does not depend on network parameters (as we show below). In this way, our model suggests capacity limits are a necessary tradeoff for the flexibility of working memory.

The model makes specific predictions about how multiple memories should impact neural activity. First, increasing the number of stimuli provided to the sensory sub-networks increases the overall average firing rate in the random network, saturating at the capacity limit of the network (*∼*3-4 items; Fig. 4A). This is consistent with experimental observations of gross activity levels in prefrontal and parietal cortex; both BOLD and evoked potentials increase as the number of items to be remembered is increased, saturating at an individual’s capacity limit [Curtis and D’Esposito, 2003; Vogel and Machizawa, 2004].

**Figure 4.**
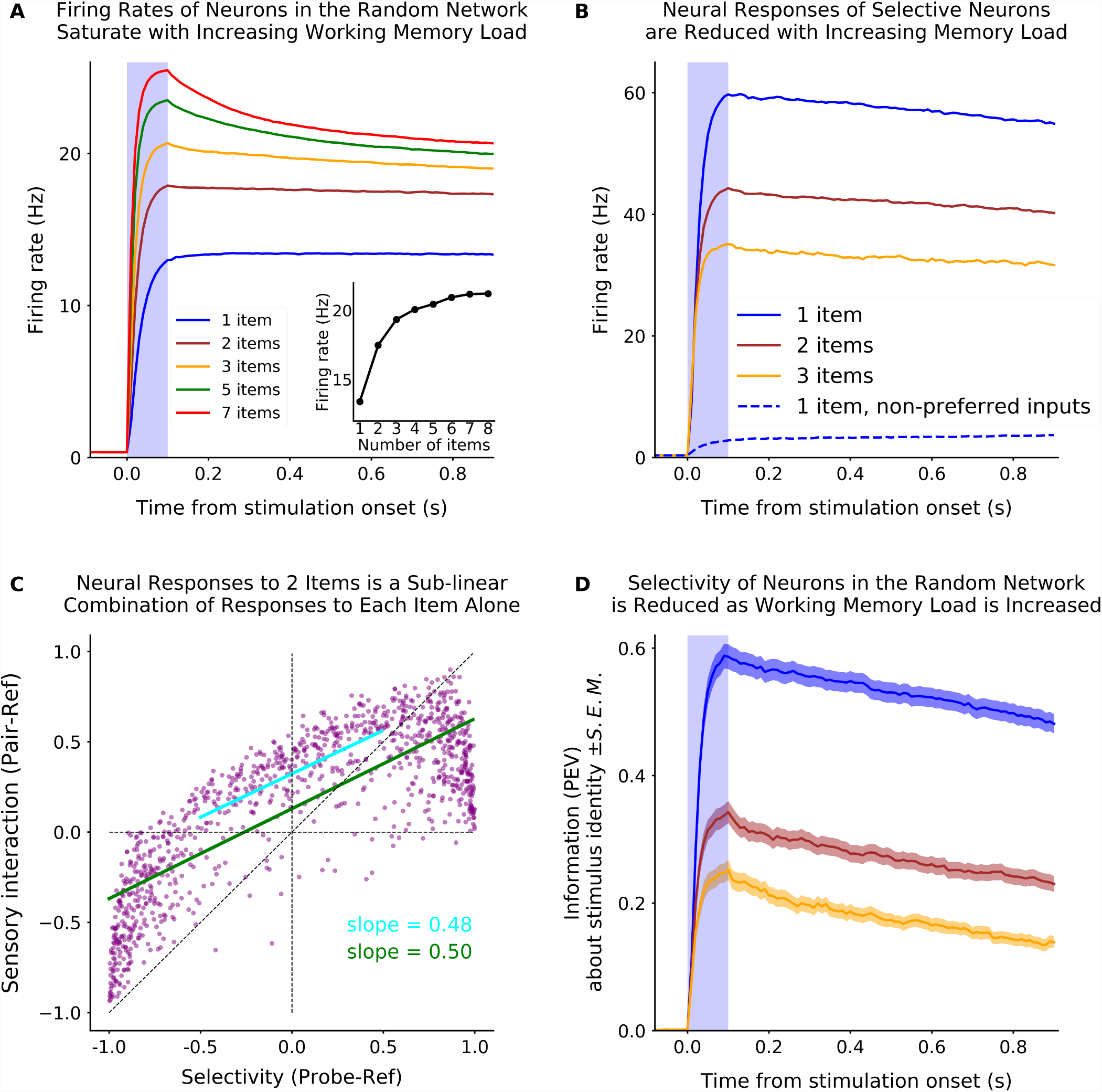
The effect of working memory load on neural responses. **(A)** The average firing rate of neurons in the random network increases with memory load, and saturates at the capacity limit of the network. **Inset:** The mean firing rate during the second half of the delay period as a function of initial load (‘number of items’). **(B)** The firing rate of selective neurons in the random network is reduced by the addition of inputs to other sub-networks. Selective neurons (N=71, 6.9% of the random network) were classified as those with a response to a preferred input that was at least 40 Hz greater than the response to other, non-preferred, inputs into other sensory networks (preferred is blue, solid; non-preferred is blue, dashed; see Methods for details). The response to a preferred stimulus (blue) is reduced when it is presented with one or two of the other non-preferred items (brown and yellow, respectively). **(C)** Divisive-normalization-like regularization of neural response is observed across the entire random network. This is shown by the response of neurons in the random network to two inputs in two sub-networks (y-axis, ‘sensory interaction’ is measured as response to the ‘pair’ of both the ‘probe’ and the ‘ref’ inputs) as a function of the response to one input alone (‘selectivity’, the ‘probe’ input, here provided into sub-network 2). All responses were relative to the response to the ‘reference’ input (here, an input into sub-network 1). A linear fit to the full distribution (green) or the central tendency (blue) shows a positive y-intercept (consistent with **A**) and a slope of 0.5, indicating the response to the pair of inputs is an even mixture of the two stimulus inputs alone. **(D)** The selectivity of neurons decreases with working memory load. The information about the identity of an input into one sensory sub-network decreases as memory load is increased (colored lines as in **B**). Information was measured as the percent of variance in the firing rate of neurons in the random network that could be explained by input identity (see Methods for details).

Second, the model predicts that the response of selective neurons in the random network will be reduced when multiple memories are maintained (Fig. 4B). This is consistent with experimental observations, which have shown divisive-normalization-like regularization of mnemonic responses in single neurons and across the population [Buschman et al., 2011; Sprague et al., 2014]. Our model provides a potential circuit mechanism for such divisive-normalization-like regularization, suggesting it is the result of balanced excitation/inhibition between networks. Neurons in the random network that respond to a single stimulus are more likely to be inhibited than excited by a second stimulus (with a ratio of 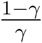). This results in an overall reduction in the response of these neurons (Fig. 4B).

This effect can be seen across the population. Figure 4C shows the relative response of neurons in the random network to two stimuli either presented separately (x-axis) or together (y-axis). As noted above, the overall average response increases with the addition of a second item (y-intercept of linear fit is 0.13 when fitting all neuron responses; 0.32 when restricting to central portion without extrema). However, the response to the pair of stimuli was not a summation of the response to each stimulus alone; rather it was a sublinear mixture (slope is *∼* 0.50 for both the full and central data). This is consistent with electrophysiological results during perception [Reynolds et al., 1999] and working memory [Buschman et al., 2011; Sprague et al., 2014]. It is also consistent with divisive normalization models that predict the response to two stimuli should be the average of the response to each stimulus alone (resulting in a slope of 0.5). Divisive normalization has been observed in many cognitive domains [Carandini and Heeger, 1994, 2012; Heeger, 1992]; our results demonstrate excitation/inhibition balance within a convergent, non-linear, network could provide one potential circuit mechanism for these effects.

The divisive-normalization-like regularization of neural responses in the random network reduces the amount of information carried about a stimulus in memory (Fig. 4D). As noted above, neurons in the random network show selectivity for the identity of stimulus inputs into a sensory sub-network, allowing them to carry information about the item in memory (Fig. 4D, blue). The addition of other items into memory reduces this selectivity, leading to a reduction in information (Fig. 4D, brown and orange for 2 and 3 items, respectively). This is consistent with electrophysiology and imaging results showing a decrease in information with increasing memory load [Buschman et al., 2011; Sprague et al., 2014].

Interference between memory representations also provides a simple mechanistic explanation for the observed decay in recall precision as a function of load [Bays et al., 2009; Bays and Husain, 2008; Ma et al., 2014; Wilken and Ma, 2004]. In our model, the memory representations drift over time, due to the accumulation of noise from Poisson variability in neural spiking [Burak and Fiete, 2012]. Greater interference in the random network leads to weaker feedback, allowing noise to have a greater impact and causing increased drift in the representations (as seen in Fig. 3C-F). In addition, reducing overall neural activity (as a result of the excitation/inhibition balance), reduces the ability to accurately decode the memory representation [Bays, 2014, 2015].

Given the suggested role of interference in memory performance, our model makes the prediction that reducing interference between memories should improve working memory performance and accuracy. Indeed, how much the representations of a pair of stimuli overlapped in the random network strongly determined the ability to accurately maintain both memories. As seen in Fig. 5A memory performance was impaired for pairs of memories with less correlated representations, reflecting greater interference between the memories. Conversely, more correlated memories were better remembered (Fig. 5A and S3). Surprisingly, memory representations were not static, but evolved over time in a way that reduced interference between memories (Fig. 5B). Both of these predictions are testable hypotheses that could be addressed with future electrophysiology or imaging experiments.

**Figure 5.**
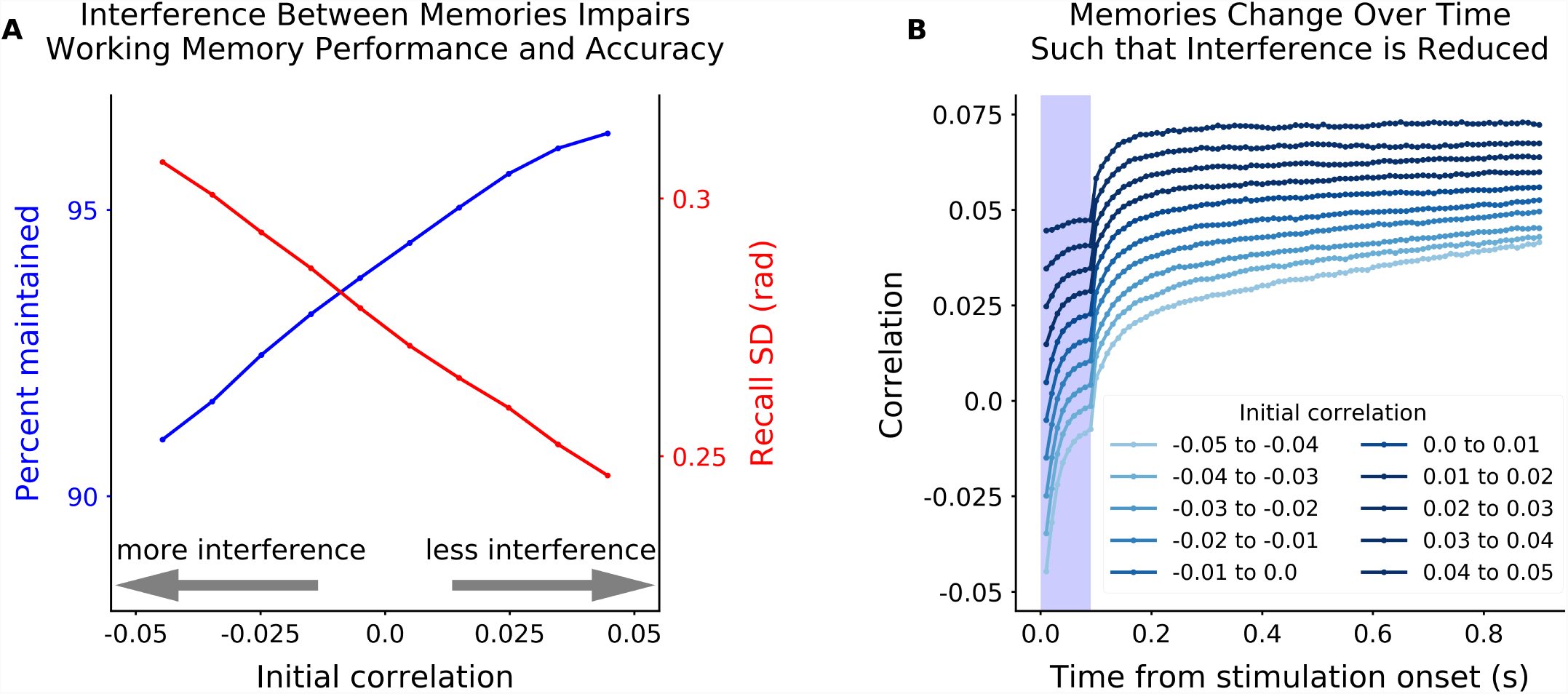
Interference between two inputs reduces performance and accuracy. **(A)** Performance (blue, as in Fig. 3A) increased as two inputs into two sensory sub-networks are more correlated in the random network. Correlation was measured as the dot product of the vector of random network responses to two inputs into two different sensory sub-networks. Random network responses were estimated by projecting the sensory input through the feedforward weight matrix. An increase in correlation reflects increasing overlap of each memory’s excitatory/inhibitory projections into the random network. Therefore, an increase in correlation reduces interference between memories and improves working memory performance. This was also true for memory accuracy, even when both memories were maintained (red, as in Fig. 3E). **(B)** Memory representations change over time in a way that increases correlation and, thus, reduces interference. Correlation was computed as in **A**, but using the maximum likelihood estimate of memory representations from the sensory network to estimate correlation (instead of the sensory input directly).

### Stable and dynamic encoding of memories

The model captures several more key electrophysiological findings related to working memory. First, we observe persistent mnemonic activity, consistent with electrophysiological results in both monkeys and humans [Funahashi et al., 1989; Fuster, 1973; Rainer and Miller, 2002; Vogel and Machizawa, 2004]. This is true, even on the first trial of being exposed to a stimulus, as observed in monkeys [Constantinidis and Klingberg, 2016]. Second, working memory activity is distributed across multiple networks, reflecting the distributed nature of working memory representations in the brain [Christophel et al., 2017]. Third, as noted above, the random nature of connections in our model yields high-dimensional, mixed-selective, representations in the random network; as has been seen in prefrontal cortex [Fusi et al., 2016; Rigotti et al., 2013]. Finally, as we show next, our model shows the same combination of stable and dynamic representations seen in neural data [Barak et al., 2010; Jun et al., 2010; Murray et al., 2016; Rainer and Miller, 2002; Stokes et al., 2017, 2013].

Recent work has highlighted the dynamic nature of working memory representations, suggesting that representations are not stable during a trial. In particular, neural representations early in the trial are not well correlated with representations later in a trial (e.g. stimulus presentation vs. memory delay, [Stokes et al., 2013]). This argues against the classic view of a static mnemonic representation. In contrast, Murray et al [2016] suggested that much of these dynamics are orthogonal to the space in which memories are represented, such that although responses are dynamic, their encoding is stable.

Our model can capture both effects. Figure 6A shows the cross-correlation across time of neurons in the random network responding to a single stimulus. Consistent with experimental observations, representations early in the trial are not well correlated with later representations (Fig. 6B). However, representations within the memory delay are relatively more stable. To account for the difference in representation during stimulus presentation, a direct projection from the input into the random network was added. In addition, weak recurrent activity within the random network was added to increase dynamics throughout the trial (see Methods for details). These biologically-plausible changes did not perturb memory performance (Fig. S5).

**Figure 6.**
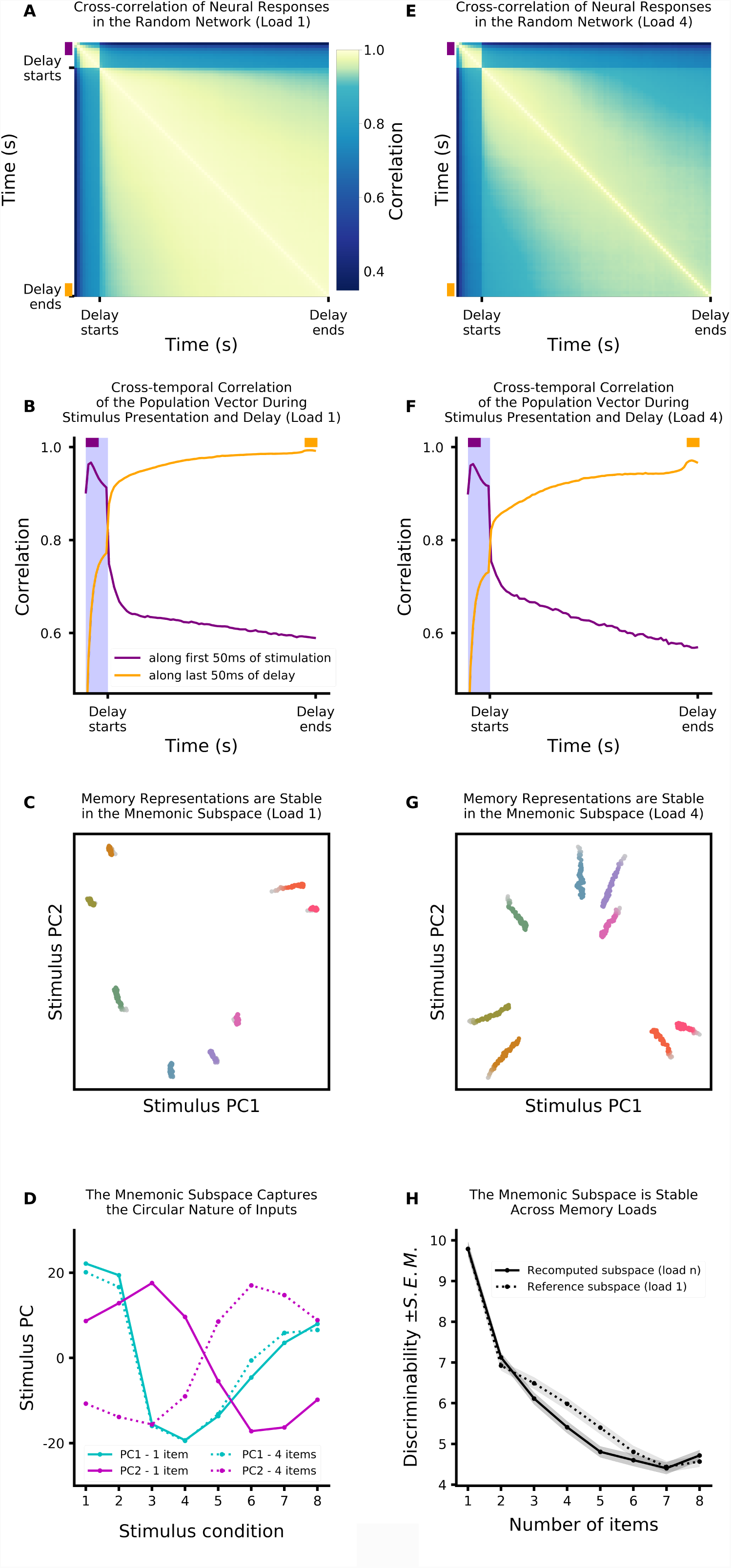
Neural dynamics are orthogonal to the mnemonic subspace. As noted in the main text, these simulations used a network with weak direct sensory input into the random network and weak recurrence within the random network (see Methods for details). **(A)** Cross-correlation of neural activity in the random network. As has been observed experimentally, correlation is low between stimulation and delay time periods, reflecting dynamic changes in the representation of memory. Dynamics during the delay period also reduce correlation across time. This is not due to forgetting: in these simulations the memory is always maintained. Correlation (color axis) is measured between the vector of firing rates in the random network for two different timepoints (along x and y axes). The color axis was chosen to be non-linear to highlight the differences between the stimulation and the delay periods. **(B)** Slices of the matrix represented in **A**: correlation of population state from the first 50ms of the stimulus period (purple) and the last 50ms of the delay period (orange) against all other times. Correlation is highest for nearby time periods, reflecting dynamics in memory representations. **(C)** Neural activity is dynamic, but memory encoding is stable. Here the response of the random network population is projected onto the mnemonic subspace (defined by the first two principal components of time-averaged activity, see Methods for details). Each trace corresponds to the response to a different input into sensory sub-network 1, shown over time (from lighter to darker colors). **(D)** Mnemonic subspace is defined by two orthogonal, quasi-sinusoidal representation of inputs, capturing the circular nature of sensory sub-networks. **(E-G)** Same as **A-C** but for a load of 4. Only includes simulations for which the memory in sensory sub-network 1 was maintained (other three memories can be forgotten). **(H)** The mnemonic subspace is stable across working memory load. Decodability of memory (here measured as discriminability, d’, between inputs, see Methods for details) is similar regardless of whether mnemonic subspace was defined for a single input (dashed line) or specific for each individual load (solid line). Decodability was not significantly different across subspaces for load 2, 3, 6, 7 and 8 (two-sample Wald test gives *p* = 0.22, *p* = 0.063, *p* = 0.32, *p* = 0.87 and *p* = 0.42, respectively). For load 4 and 5, decodability was better for the single input subspace (*p* = 0.0076 and *p* = 0.0042 respectively). In general, as expected, decodability is reduced with load (*p <* 0.001).

However, despite these dynamics, the subspace in which a memory is represented is stable. Following the approach from Murray et al. [2016], we estimated the ‘mnemonic subspace’ of the random network by finding the first two dimensions that explained the most variance of the time-averaged response to eight different stimulus inputs (see Methods for details and Fig. S6B). Projecting the response of the random network over time revealed the representation was stable (Fig. 6C; note that this subspace is not constrained to minimize temporal variability). Furthermore, despite the randomness of connections, the representational subspace of the stimuli was a mixture of two orthogonal sinusoids, reflecting the circular nature of inputs into the sensory network (Fig. 6D, solid lines). Remarkably, this matches the stable subspace found in prefrontal cortex [Murray et al., 2016]. Together, these results show how the low (two) dimensional space of each stimulus network is embedded in the higher dimensional space of the random network.

Previous experimental work has largely focused on the representation of single items in working memory. Our model makes several predictions for how these representations should change when the number of items in working memory is increased. First, we found that increasing working memory load decreased the cross-correlation over time (Fig. 6E). This is due to a decrease in signal-to-noise ratio with load as well as interactions between memory items, as discussed above. Second, we found an increase in the drift of representations through the mnemonic subspace (Fig. 6G). These simulations only include those where the memory was successfully maintained, therefore this does not reflect drift toward a null state (where the memory is forgotten). Instead, it reflects the weakening of memory representations due to interference. This reduces the discriminability of memories (Fig. 6H, solid line), impairing working memory performance.

Despite the addition of multiple memories, the mnemonic subspace for decoding the memory from one sub-network did not change. The subspace for the first item, when it was remembered with 3 other items, was a combination of two orthogonal sinusoids (Fig. 6D, dashed lines). Furthermore, we found that the subspace that encoded a stimulus in a sensory sub-network was stable across memory loads. This means one could use the decoding subspace for a memory when it was presented alone to decode that same item when it was remembered with other stimuli. Indeed, decoding performance was equal, or a bit better, when using the mnemonic subspace from a single item compared to the subspace defined at each load (Fig. 6H and Figure S7B). This is critical for being able to use the representations within the random network – as the subspace doesn’t change, any learning done at one memory load would generalize to other loads.

These results also highlight how the mnemonic subspaces for each sensory network are nearly orthogonal to one another when projected into the high-dimensional space of the random network. This can also account for how dynamics in the random network do not impact memory representations – as long as the dynamics are orthogonal to the mnemonic subspace of the sensory network, they will not alter memories. Indeed, the structure embedded in the sensory network (through the center-surround connectivity) can help to constrain dynamics within the random network to irrelevant dimensions.

### The network is robust to changes in architecture, parameters and connectivity

Our model demonstrates how random connections can support the flexible maintenance of memories. To highlight the flexibility of the model with respect to the architecture of sensory inputs, we replaced four of the ring-like sensory sub-networks with line networks, modeling a one-dimensional psychometric spaces such as brightness or spatial frequency (Fig. S8A). Without changing any parameters, the network was able to maintain inputs into both circular and linear spaces (Fig. S8B). This highlights the strength of the model – it does not rely on finely tuned attractors for each type of information that must be maintained. In order to maintain ease of comparison with the extant theoretical and experimental literature, we focused our simulations on circular sensory sub-networks.

As noted above, network parameters were set to maximize the total number of remembered stimuli and minimize the number of spurious memories, at all loads (see Methods for details). Maintenance of working memory representations in our model relies on sufficient recurrent activity between the sensory and random networks. This constrains both the weights and the pattern of connectivity between the sensory and random networks.

First, to maintain a representation, feedforward and feedback weights between the sensory and random networks must be set such that a single action potential in the sensory network leads to roughly one action potential in the random network on average (and vice versa). This is important to sustain representations between the networks. However, the exact value can be partially relaxed without loss of functionality. First, what matters is the product of feedforward and feedback weights. Thus, several possible values of feedforward/feedback weights give nearly identical network performance as an increase in feedforward weight can be compensated by a decrease in feedback weight (or vice versa, Fig. 7A, inset). Second, the product can also be changed up to 5% without significant loss of function (Fig. 7A). Beyond this range, there is either insufficient drive to sustain activity (weights are too low), leading to loss of representations (Fig. 7A, below *-* 5%); or there is unchecked amplification of activity (weights are too high), leading to spurious representations throughout the network (Fig. 7A, above +5%).

**Figure 7.**
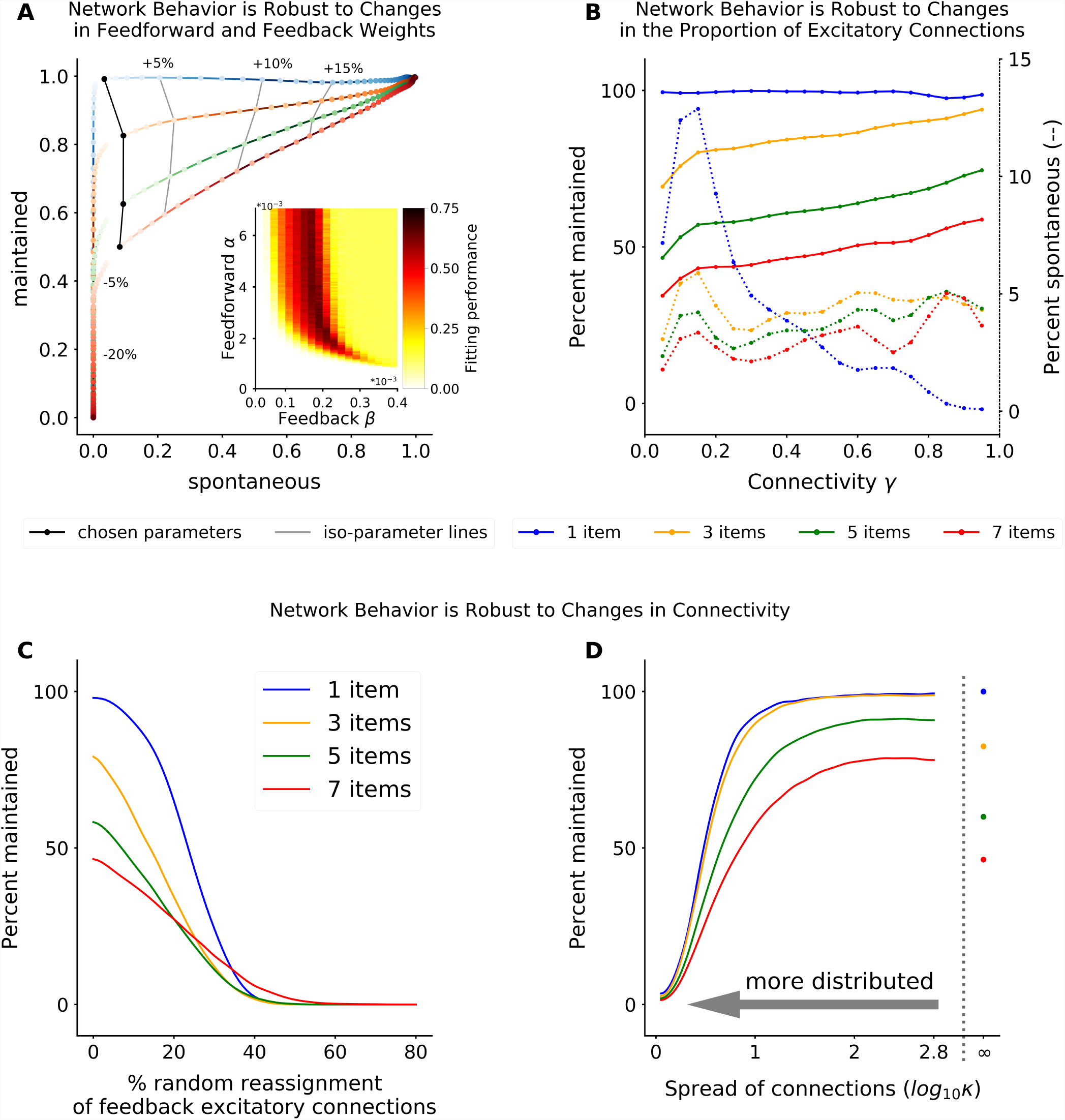
The network is robust to changes in parameters. **(A)** ROC plot showing how changing feedforward/feedback weights influences network performance. The probability of correctly maintaining a memory (hit rate; y-axis) and the probability of creating a spontaneous (spurious) memory in non-stimulated sensory sub-networks (false alarm; x-axis) varies as the product of the feedforward and feedback weights are changed relative to optimal values (darker colors move away from optimal value). Isolines show network performace across memory load when the product of feed-forward and feedback weights is changed by 5%, 10%, and 15%. Performance decreases with memory load (colored lines) for all parameter values. Here *γ* = 0.35. **Inset**: Network performance (color axis) as a function of feedforward and feedback weights. Network performance was taken as the percent of trials with successfully remembered memories minus the percent of trials with spontaneous memories (see Methods for details). There is a large space of well-performing parameters, although network performance most critically depends on the product of feedforward and feedback weights. Here *γ* = 0.35. **(B)** Optimal feedforward and feedback weights can be found for a broad range of *γ* values that maximize the percent of inputs maintained (solid lines) and minimize the number of spontaneous memories (dashed line). Again, memory performance is decreased with load (colored lines), for all parameters. **(C and D)** Network behavior is robust to changes in connectivity. The percent of remembered inputs (y-axis) for different memory loads (colored lines) decreases as connections between random and sensory networks are either (**C**) randomly re-assigned (breaking symmetry) or (**D**) locally redistributed. As seen in **C**, the network is partially robust to breaking symmetrical connections between sensory and random networks. However, the network is much more robust when these connections are locally distributed within a sensory network (**D**). See Methods for details and Fig. S9 for examples of redistribution for different values of *κ*. Due to bit depth constrains, Von Mises are only calculated for log_10_ *κ <* 2.8. The symbol is used for the limit of no redistribution, thus corresponding to a Dirac delta function.

Similarly, the fraction of connectivity between neurons in the random network and the sensory networks (*γ*) can be significantly changed without any qualitative change in network behavior (assuming the connection weights are adjusted to meet the above criterion). Fraction of connectivity ranging from approximately 30% to 95% show similar flexibility and capacity limits (Figure 7B). Here we chose *γ* = 0.35 to maximize maintenance, minimize spurious memories, and minimize connections (thereby minimizing structural costs).

The final constraint on connectivity is that the neurons between the sensory and random network are reciprocally connected. This is consistent with wiring in the brain – neurons are more likely to be reciprocally connected than expected by chance (see below for biological mechanisms, [Holmgren et al., 2003; Ko et al., 2011; Markram et al., 1997; Song et al., 2005]). However, as with the other parameters in the model, the reciprocal nature of connections can be relaxed significantly without degrading network performance. To demonstrate this, we randomly reassigned a proportion of the feedback excitatory connections from one sensory neuron to another (even in a different sensory network; all model parameters were left unchanged). The model was robust to at least 10% of excitatory connections being randomly switched with an inhibitory connection (to any sensory sub-network; Fig. 7C). However, biological connection errors are more likely to be spatially localized and so we tested whether the network is also robust to distributing excitatory feedback connections to nearby neurons within a sensory network. The network is robust to these changes, able to sustain memories without a significant decrease in performance, even when distributing half of the connection weight over more than 45*°* away from its initial position in the sensory sub-network (i.e. *κ ≥* 1, Fig. 7D and S9). In fact, when connections were distributed over a small range of nearby neurons (*κ ∼* 2), performance of the network was improved due to the fact that smoothing feedback connections reduced the impact of stochasticity in neural activity. This suggests the local distribution of connections observed in the brain may actually be advantageous.

Altogether, these results demonstrate that our network architecture is fairly robust to changes in parameters. Importantly, for all parameters tested the network was flexible and had a limited capacity.

## Discussion

We present a simple model of working memory that relies on random and recurrent connectivity between a structured sensory network and an unstructured random network. Despite its simplicity, the model captures several key behavioral and neurophysiological findings. In particular, the model demonstrates how random, convergent connections can support the flexibility of working memory. However, this flexibility comes at the cost of a limited capacity. Interference between multiple memories leads to divisive-normalization-like regularization of responses which, in turn, leads to an increase in memory error and, eventually, to a failure to maintain some memories (forgetting). Additionally, the model explains 1) how working memory representations can be both static and dynamic; 2) observations that working memory is distributed across the cortex; 3) both the general increase in neural activity with working memory load as well as regularization at the single neuron level; and 4) the conjunctive stimulus tuning properties of neurons in associative (e.g. prefrontal) cortex.

### Biological Plausibility

The sensory networks in our model are intended to capture two key aspects of generic sensory cortex. First, neurons receive tuned inputs, giving rise to selectivity in their responses [Funahashi et al., 1989; Mendoza-Halliday et al., 2014; Zaksas and Pasternak, 2006]. Second, there is a center-surround architecture with respect to tuning. Pyramidal neurons with similar selectivity are more likely to be connected to one another [Holmgren et al., 2003; Ko et al., 2011; Markram et al., 1997; Song et al., 2005] while inhibitory interneurons spread inhibition across the population (likely through parvalbumin-positive inhibitory interneurons [Atallah et al., 2012; Packer and Yuste, 2011; Wilson et al., 2012]). Such structure is thought to emerge through unsupervised learning mechanisms, such as Hebbian plasticity [Hebb, 1949]. In this way, it reflects the statistics of the world, embedding knowledge about how stimuli relate to one another. This is critical to the model, as it constrains the activity in the random network such that it stays within a biologically meaningful subspace, allowing it to be easily interpreted for behavior.

In contrast to the sensory network, the random network in our model is unstructured. Similar to reservoir pool computing approaches in machine learning [Jaeger, 2002], the random connections to the random network create a high dimensional space where any type of information can be represented. This is what endows our model with its flexibility. Experimental results have suggested that associative regions, such as prefrontal cortex or the hippocampus, may have such high-dimensional representations [Fusi et al., 2016; McKenzie et al., 2014; Rigotti et al., 2013] and therefore could be a possible physical instantiation of the random network. In particular, prefrontal cortex is known to have strong bi-directional connections throughout sensory cortex, making it particularly well-suited to play the functional role of the random network [Duncan, 2001; Miller and Cohen, 2001]. Indeed, lesioning prefrontal cortex severely impairs working memory performance [Jacobsen et al., 1936; Levy and Goldman-Rakic, 1999; Petrides, 1995].

One molecular mechanism that could support reciprocal random connectivity are protocadherins, a subclass of cell adhesion molecules. Protocadherins undergo a recombination process such that each neuron expresses a unique set of these cell adhesion molecules [De Wit and Ghosh, 2016]. Neurons with overlapping expression profiles of protocadherins are more likely to make synaptic connections [Kostadinov and Sanes, 2015]. Generally, this is important for functioning of neural networks: randomization of initial connections is critical for effective learning in artificial neural networks. Here, we propose that this randomization facilitates the formation of a reservoir pool in the random network, providing the architecture for the flexibility of working memory.

Our network uses balanced excitation and inhibition to constrain neural activity to a stable regime. This is consistent with recent experimental work that has found excitation and inhibition are tightly balanced within the cortex [Mariño et al., 2005; Vogels and Abbott, 2005; Vogels et al., 2011]. Although long-range connections, such as modeled between the sensory and random networks, are largely excitatory in the brain, feed-forward inhibition within the target region can ensure these projections are effectively excitation/inhibition balanced.

Although the model is biologically plausible, it is intentionally simple. Several simplifying assumptions were made: neurons do not obey Dale’s law, there are no cell types, and there are no interactions between sensory sub-networks. Here, our goal was to demonstrate the explanatory power of the simplest network. This highlights the computational power of random, convergent, connections, showing how they can account for the flexibility of working memory, as well as its limited capacity.

However, a few aspects of working memory representations remain unexplained by our model. For example, recent results suggest short-term memory relies, at least in part, on short-term synaptic changes [Mongillo et al., 2008; Postle, 2017; Stokes, 2015; Stokes et al., 2017] or oscillations [Lundqvist et al., 2016]. The simplified biophysics of our neurons do not capture these effects; future work will investigate whether extending the model can explain these additional findings.

### Model Predictions

In addition to capturing existing behavioral and electrophysiological results, the model makes several testable hypotheses. First, the model predicts that memory performance should be a function of how strongly neural representations interfere with one another. As we show above, interference between two items can be estimated from the correlation between representations when each item is presented alone. Our model predicts increasing interference should reduce behavioral performance. Furthermore, it suggests that memories should drift over time in a way that reduces interference between memories.

Second, the model predicts that disrupting random connectivity in the cortex should disrupt working memory performance. Indeed, recent results suggest this may be true – mice with reduced diversity in protocadherins have impairments in sensory integration and short-term memory [Yamagishi et al., 2018]. At the single neuron level, the model would predict a reduction in the diversity of conjunctive representations in associative cortex when protocadherin diversity is impaired.

Finally, the model predicts that maintaining a stimulus presented at one location should increase neural activity in other locations in sensory cortex. This activity should be consistent from trial to trial, but unstructured with respect to the content held in memory. Although, in vivo, lateral projections within sensory cortex (which are not included in our model) may provide some structure to the distributed activity or act to reduce noise.

### Filtering of complex top-down projections by local structure

Our results show how complex representations in higher order cortical regions can be filtered by the structure of lower cortical regions. The random network represents multiple items in a high dimensional space. However, because of the structure within each sensory sub-network, only the component relevant to that sensory sub-network affected representations. In this way, the complex representations observed in higher-order brain regions (e.g. PFC, Rigotti et al. [2013]) can influence activity in lower-order, sensory regions. While our model shows how this can be used to hold items in working memory, this same concept can apply to other forms of top-down control, such as cognitive control or attention.

The necessity for local structure in filtering top-down projections is not due to the random connections used in our current model. Any degree of convergence along the cortical hierarchy will mean that the specificity of representations in higher order cortex must be lower than that of lower order cortex. Therefore, in order for top-down projections to act at the specificity of lower cortical regions, additional constraints must be added. Our model shows structure within the lower cortical region could provide these constraints.

Finally, filtering by structure in lower cortical regions may also facilitate learning of top-down control signals. Learning in high-dimensional spaces is an unavoidably difficult problem; by filtering the top-down signal through the local network, only the relevant components will impact behavior, limiting the effective dimensionality in which the control signal must be learned.

### The trade-off between flexibility and stability

The random architecture of our model supports its flexibility. However, this comes at the cost of a limited capacity when multiple memories interfere with one another. This interference could be reduced by forming independent networks for each type of information one wants to maintain in working memory. While this comes at the loss of flexibility, it would form a more stable representation. There is some evidence for such network specialization in the brain: spatial location, such as heading direction, may be represented in specialized ring attractors ([Kim et al., 2017; Seelig and Jayaraman, 2015]).

This trade-off between flexibility and stability exists for all systems in the brain. Behaviors can be represented in specialized networks, such as when learning a motor skill (e.g. walking or chewing gum). Such independent representations are robust to noise and reduce interference with other behaviors. However, they require extended periods of learning (either across evolution or during an organism’s lifetime). In contrast, our network architecture provides the structure needed for generalized behavior. Indeed, the flexible representations encoded in our model are reminiscent of the selectivity seen in prefrontal cortex [Fusi et al., 2016; Rigotti et al., 2013] and as predicted by flexible models of cognitive control [Duncan, 2001; Miller and Cohen, 2001]. Our results show how such dynamic, adaptive representations can interact with the knowledge embedded in structured networks to constrain representations to be behaviorally relevant, supporting cognitive control of behavior.

## Acknowledgements

The authors are grateful to Sarah Henrickson, Adam Charles, Alex Libby, Camden MacDowell, Sebastian Musslick, and Matt Panichello for discussions and feedback on the manuscript. This work was supported by NIMH R56MH115042 and ONR N000141410681 to TJB.

## Author Contributions

TJB initially conceived of the model; FB and TJB designed the final model, implemented the model, discussed the results and wrote the manuscript.

## Declaration of Interests

The authors declare no competing interests.

## Methods

### Computational model

We consider a two-layer network of Poisson spiking neurons. Model parameters were adapted from [Burak and Fiete, 2012]. Each neuron *i* generates Poisson spikes based on its time-varying firing rate *r*_*i*_(*t*). The firing rate of neuron *i* is a non-linear function of the weighted sum of all pre-synaptic inputs:

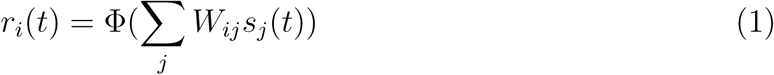

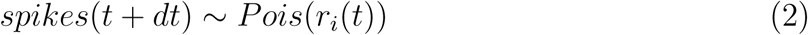

where *W*_*ij*_ is the synaptic strength from pre-synaptic neuron *j* to post-synaptic neuron *i*; *s*_*j*_(*t*) is the synaptic activation of pre-synaptic neuron *j*; and Φ is a baseline-shifted hyperbolic tangent: *τ* Φ(*g*) = 0.4(1 + tanh(0.4*g -* 3)). The synaptic activation produced by neuron *j* is computed by convolving its spike train with an exponential function:

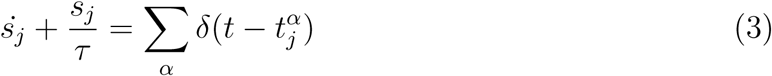

where 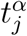 are the spike times of neuron *j*. For simplicity, we choose the synaptic time constant *τ* = 10*ms* to be equal for all synapses. As noted in the main text, we are using a simplified model of neural activity; neglecting cell-types, the existence of refractory periods, and bursts of neural activity.

Previous work has shown the variability in spiking activity plays a significant role in the diffusion of memories over time ([Burak and Fiete, 2012]), which motivated our choice of using spiking neurons instead of a rate model. All variability in spiking arises from the inhomogeneous Poisson process used to generate spike events (i.e. no noise was added).

The model consists of two interacting layers of neurons: a ‘sensory’ network and a ‘random’ network (Fig. 1A). The sensory network consists of 8 independent ring-like sub-networks. These sub-networks are each composed of *N*_*sensory*_ = 512 neurons and mimic simplified sensory networks, with neurons around the ring corresponding to specific values of a continuously encoded feature, such as shape, orientation, or color (here graphically represented as color). Every neuron *i* has an associated angle *θ*_*i*_ = 2*πi/N*_*sensory*_. Consistent with biological observations [Funahashi et al., 1989; Kiyonaga and Egner, 2016; Kuffler, 1953], connections within a sensory sub-network have a center-surround structure (Fig. 1A, inset). The synaptic weight between any pair of neurons is rotationally invariant, with nearby excitation and surround inhibition. The synaptic weight between any pair of neurons *i, j* within a sensory sub-network depends only on *θ* = *θ*_*i*_ *-θ*_*j*_ through:

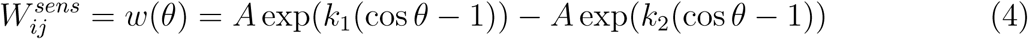

where *k*_1_ = 1 is the width of the excitation kernel, *k*_2_ = 0.25 is the width of the suppression kernel, and *A* = 2 is the amplitude. Self-excitation is set to 0 (*w*(0) = 0).

Different sub-networks reflect independent sensory inputs (either stimuli at different locations in space or different sensory features or modalities). In the current model the sensory sub-networks are independent (unconnected), although adding weak inhibition between sub-networks did not qualitatively change our results.

The random network is composed of *N*_*rand*_ = 1024 neurons randomly connected to each of the 8 simulated sensory sub-networks (four-fold compression). As there is a single random network, all sensory sub-networks converge onto it through a random feed-forward connectivity matrix (*W* ^*FF*^). Each neuron in the random network then feeds back into the same random subset of neurons in the sensory network that provided excitatory inputs. In other words, the connections were bi-directional and therefore, the feedback connectivity matrix (*W* ^*FB*^) from the random network to the sensory network is the transpose of the feedforward connectivity matrix, with distinct weight values (i.e. 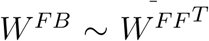). The likelihood of an excitatory connection between any pair of neurons across the sensory and random networks was defined by *γ* (default is 0.35). Therefore, the number of inter-network excitatory connections (*N*_*exc*_) for any given neuron followed a binomial distribution (with *p* = *γ*). The strength of the excitatory feedforward and feedback connections were defined by the parameters *α* and *β*, respectively (how these were chosen is detailed below).

Connection weights between the sensory and random networks were balanced in two ways. First, in order to balance the total excitatory drive across neurons, feedforward (or feedback) weights were scaled by *N*_*exc*_. Second, the connections between the sensory and random networks were balanced such that individual neurons receive an equal amount of excitatory and inhibitory drive (i.e. 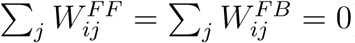). To achieve this, an equal inhibitory weight was applied to all inputs for each neuron 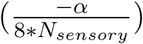. This method of balancing is intended to reflect broad inhibition in the target network due to local inhibitory interneurons.

Thus, after balancing, excitatory feedforward connections from the sensory to random network neuron *i* had weight 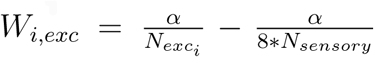 while inhibitory feedforward connections had weight 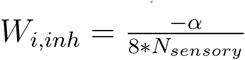. Similarly, neuron *j* in the sensory network will receive excitatory feedback connections with weight 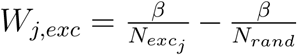 and inhibitory feedback connections with weight 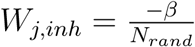.

### Presentation of stimuli to the sensory network

Sensory stimuli were provided as synaptic drive (*s*_*ext*_) to the sensory sub-networks:

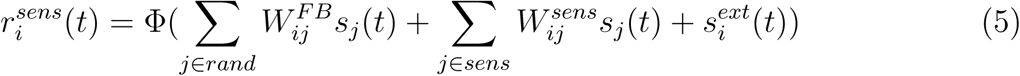

All inputs were presented for 100 ms, indicated by the blue shaded region in all figures. Inputs were Gaussian, with the center of the input (*µ*) chosen randomly for each sensory sub-network. The width of the input (*σ*) was defined as a fraction of the total number of neurons in the sensory sub-network, 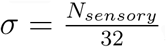, which was 16 neurons for the presented network. Inputs beyond three *σ* were set to 0. Therefore, the sensory input to neuron *i* was

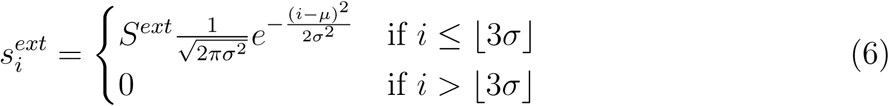

where *S*^*ext*^ was the strength of external sensory input (here, *S*^*ext*^ = 10).

### Adding direct sensory input and recurrence within random network

The baseline model used for most analyses is intentionally simple. The activity of neurons in the sensory network was given by Equation 5 and the activity of neurons in the random network was:

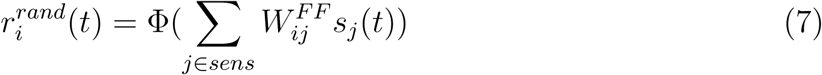

However, to investigate the dynamics of activity within the network, two biologically-motivated connections were added (only for results presented in Fig. 6). First, inputs were projected directly into the random network (in addition to inputs into the sensory network). A random weight matrix, 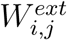 defined the feedforward connectivity of sensory inputs into the random network. Similar to the weight matrix from the sensory network, the likelihood of an excitatory connection from any given input was *γ*^*ext*^ (default is 0.2) with weight proportional to *α*^*ext*^. As with other connections in the network, inhibitory connections were added to all neurons in the random network in order to balance excitation and inhibition. Second, recurrence within the random network was added. Again, a random weight matrix was constructed 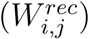 with a likelihood of excitatory connection defined by *γ*^*rec*^ (default 0.2) and with weight proportional to *α*^*rec*^. As with feedforward and feedback connections, these connections were balanced such that overall excitatory drive was constant across neurons and excitation and inhibition were balanced within a neuron.

Therefore, in the network used to investigate dynamics (Fig. 6), the activity of neurons in the random network was:

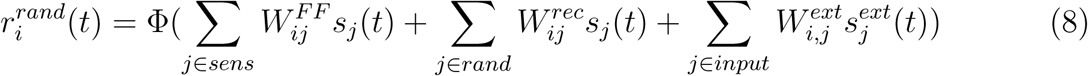

### Performance of the network

Network performance or ‘percent maintained’, in Figures 3, 5 and 7 was defined by the proportion of successfully maintained memories with respect to the number of initial inputs to sensory sub-networks. A memory was ‘maintained’ if the norm of the activity vector for a given sub-network exceeded 3Hz. We did not choose a similarity metric with the initial representation of inputs, because that would not allowed for taking memory drift into account. Similarly, a sub-network carried a ‘spontaneous’ memory if it did not receive any input during stimulation but had activity that exceeded the same threshold.

### Parameter estimation of the weight matrix between the sensory network and the random network

The only constraint on network sizes is the fact that the random network is convergent (*N*_*rand*_ *<* 8 *\* N*_*sensory*_). Here we use a four-fold convergence, although changing this did not qualitatively change network behavior. A limit case would be to have both networks of the same sizes, with one-to-one, non-overlapping representation of inputs from the sensory network to the random network, but this case is not in the interest of the study.

Feedforward and feedback connection weights (*α* and *β*, respectively) were set using a grid search that optimized model performance over hundreds of simulated trials. As described in the Results, the exact values of network parameters can be partially relaxed without changing network behavior. The parameters were fit to optimize two constraints: 1) maximize the number of inputs that were maintained throughout the stimulation and the delay periods and 2) minimize the number of spontaneous memories. We chose these criteria as they satisfied the minimal requirements for working memory: maintaining items without hallucinating. The performance of a set of parameters was evaluated as the sum of the proportion of maintaining a memory minus the proportion of creating a spontaneous, summed for simulations for initial inputs number from 1 to 8 (Fig. 7A). Despite attempting to optimize for overall network performance, no parameters were found that escaped the capacity limitation.

For a broad range of *γ*, we can find *α* and *β* parameters that maintain stimulus inputs and minimize spontaneous memories (Fig. 7B). We set *γ* to 0.35 as it roughly maximized the number of maintained stimuli, minimized the number of spontaneous memories, and minimized the number of connections. The corresponding best fit for feedforward and feedback connections was *α* = 2100 and *β* = 200. This resulted in an average excitatory feedforward connection weight of 0.95 and an average excitatory feedback connection weight of 0.36. From the balance, we can compute that the inhibitory feedforward connection weight is −0.51 and the inhibitory feedback weight is −0.20. Note that, for all parameters tested, interference between inputs into the random network limits memory maintenance and accuracy – the capacity of the network is unavoidable.

For the model that included direct sensory inputs into the random network and re-currence within the random network, *α*^*exc*^ and *α*^*rec*^ were fit using a coarse grid search. As for *α* and *β*, parameters were chosen to maximize maintenance of memories while minimizing spurious memories. In addition, parameters were constrained to be positive in order to ensure transient dynamics in neural activity. The best fit parameters were *α*_*input*_ = 100 and *α*_*rec*_ = 250. Figure S5 shows that the addition of these connections does not qualitatively change the model behavior.

### Estimating memory accuracy

Building on [Bays, 2014; Pouget et al., 2000], memories were decoded from the sensory sub-networks using a maximum likelihood decoder on spiking activity in a 100 ms time window (shortening the decoding interval to 50 ms or 10 ms did not qualitatively change the results). In Figure 3C,D, and Figure S2, all simulations were taken into account, even when the memories were forgotten. In Figure 3E,F, only the simulations where all the memories were maintained were taken into account.

### Measuring correlation between memories in the random network

In Figure 5 and S3, we show how memory performance increases as the correlation between memories in the random network is increased (i.e. there is a reduction in interference). The correlation between memories was computed as the dot product of the sensory inputs projected into the random network. This projection was done by multiplying the sensory input by the feedforward connection matrix. When calculating the correlation over time, we used the maximum likelihood decoded representation as our representation in the sensory network.

### Measuring divisive-normalization-like regularization of neural responses with memory load

Neural responses are reduced with an increase in load, both in the sensory domain [Carandini and Heeger, 1994, 2012; Heeger, 1992; Heeger et al., 1996] and in working memory [Buschman et al., 2011; Sprague et al., 2014]. We tested whether similar divisive-normalization-like regularization was seen in our network model.

First, we tested whether the response of ‘selective’ neurons was reduced as memory load was increased. To this end, we simulated trials where each of three different sensory sub-networks received an input (*s*_1_, *s*_2_, or *s*_3_ into the first, second, or third sub-network, respectively). This can be conceptualized as presenting a stimulus at one of three different locations. Neurons in the random network were classified as ‘selective’ if their response to any individual input (the ‘preferred’ input) was greater than the other two inputs (‘non-preferred’ inputs). This follows the procedure typically used for electrophysiology experiments. Requiring a difference of at least 40 Hz resulted in 71 selective cells (6.9%). The response of selective neurons was then computed when the preferred input was presented alone (load 1), with either of the other two non-preferred inputs (load 2), or with all three inputs (load 3). We considered only the simulations where all inputs were maintained over the delay period. As seen in Fig. 4B, increasing memory load reduced the activity of selective neurons. Similar results were observed with selectivity thresholds of 50 Hz or 60 Hz.

To quantify the impact of memory load on all neurons, we adapted the statistic from [Reynolds et al., 1999] to measure the response of a neuron to a pair of stimuli, relative to the response to each stimulus alone (Fig. 4C). The response of all neurons in the random network were calculated for three conditions: a single input to sensory sub-network 1 (*s*_1_), a single input to sensory sub-network 2 (*s*_2_), and inputs to both sensory sub-network 1 and 2 simultaneously (*s*_1,2_). As in [Reynolds et al., 1999], responses were normalized by the maximum response for each neuron (max(*s*_1_, *s*_2_, *s*_1,2_)). Figure 4C shows the relative change in response to the two single stimuli (x-axis, *s*_2_ *- s*_1_) against the pair of stimuli (y-axis, *s*_1,2_ *- s*_1_). A linear fit was used to estimate the slope of the response. The resulting slope of 0.5 indicates the response to a pair of stimuli (*s*_1,2_) is a mixture of the two stimuli presented alone, as in [Reynolds et al., 1999].

Finally, we quantified whether increasing memory load reduced the selectivity of neurons. To this end, we calculated the response of neurons in the random network to two different inputs into the first sensory sub-network (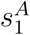 and 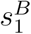). The two sensory inputs were evenly separated (offset by 180*°*). As above, neurons were classified as selective if their response to the two inputs differed by at least 40 Hz 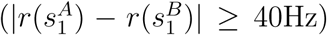. This resulted in 67 selective neurons in the random network (6.5%). The response of these selective neurons was then calculated when each input (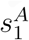 and 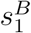) was presented with a second (or third) input into sub-network 2 (or 3), reflecting an increasing memory load. Again, we considered only the simulations where all inputs were maintained over the delay period. Following [Buschman et al., 2011], we computed the information these selective neurons carried about the identity of the stimulus in sensory sub-network 1 using the percentage of explained variance (PEV) statistic. Specifically, we computed for each selective neuron and for each time step, *η*^2^ = *SS*_*between*_*/SS*_*total*_ where 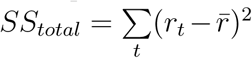 is the total squared error across trials (*t*) and 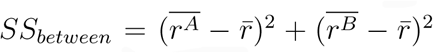 where 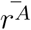 and 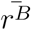 are the average response to sensory input 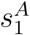 and 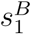, respectively. Figure 4D shows *η*^2^ decreases with increasing memory load, as seen experimentally [Buschman et al., 2011]. Similar results were seen with a threshold of 50 or 60 Hz.

### Subspace Analysis

Analysis of the stable mnemonic space (Figs. 6 as well as S4 and S6) followed the methods of [Murray et al., 2016]. To understand the subspace used for encoding, we calculated the response of the network to 8 different inputs into the first sensory sub-network 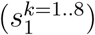. The number of stimulus conditions was chosen to match [Murray et al., 2016], corresponding to 8 angular locations evenly separated within the sub-network (i.e. spaced by 45*°*). For each input condition, 500 independent trials were simulated for the same network. We took into account only the simulations for which the memory in the first sensory sub-network is maintained after the delay period.

We assessed temporal dynamics of neural activity by computing the cross-correlation of responses across trials with the same input. Correlations were then averaged across input conditions, yielding the cross-correlation matrix in Fig. 6A. Slices of the cross-correlation are displayed in Fig. 6B to match recent findings [Murray et al., 2016; Stokes et al., 2013].

Next, we defined the mnemonic and temporal subspaces as in [Murray et al., 2016]. First, we averaged the activity of each neuron over trials. This resulted in a high-dimensional state space defined by the firing rate of *N*_*rand*_ neurons, over time *T*, for each of 8 input conditions (i.e. a *N*_*rand*_ × *T* × 8 matrix). Then, we defined the mnemonic subspace as the first two principal components of the time-averaged response (which would now be a *N*_*rand*_ × 1 × 8 matrix). Projecting the original full timecourse of neural activity onto this mnemonic subspace gave the 8 clusters of time-varying activity in Fig. 6C.

In order to analyze how memory load impacted the mnemonic subspace (Fig. 6H and Fig. S7), we repeated this process for memory loads 2 through 8. Memory load was manipulated by providing a random input into sensory sub-networks 2 through 8, in addition to the input to sub-network 1. We took into account only the simulations for which the memory in the first sensory sub-network is maintained. The memories of the other sensory sub-networks stimulated can be forgotten. This provided new mnemonic subspaces for each load.

Discriminability was computed as the distance between the neural response to different input conditions, normalized by the standard deviation of responses (*d′*). Neural responses were calculated as the average response over the delay period. The response for each trials was then projected into the mnemonic subspace defined for a stimulus presented alone (the reference subspace, load = 1) or optimized for each memory load. *d′* was calculated for each pair of inputs, along the vector connecting each cluster’s barycenter. The overall discriminability was then taken as the average *d′* across all pairs of inputs. Standard error was estimated by bootstrapping (1000 draws). Discriminability was computed within the reference (load 1) subspace and within the subspace optimized for each load. As seen in Fig. 6H, decodability was nearly equal across subspaces. For completeness, we also built a nearest centroid classifier as in [Murray et al., 2016]. Decoder accuracy was estimated by testing cross-validation performance on withheld trials, for both the reference (load 1) subspace and the subspaces optimized for each load (Fig. S7). Standard error was estimated by bootstrapping (1000 draws). This second analysis also shows that decodability was nearly equal across subspaces.

As noted above, all subspace analyses used the model with direct sensory input into the random network as well as recurrence within the random network.

### Software

All simulations were done with Python 2.7 (using numpy and scipy) and the Brian2 simulator [Goodman et al., 2014; Stimberg et al., 2013], using exact integration with time-step *dt* = 0.1*ms*. For PCA and the nearest centroid classifier, we used the scikit-learn package [Pedregosa et al., 2011].

The datasets generated from simulations and the script of the model will be provided on GitHub after finalizing the manuscript.

## Supplemental Figures

**Figure S1.**
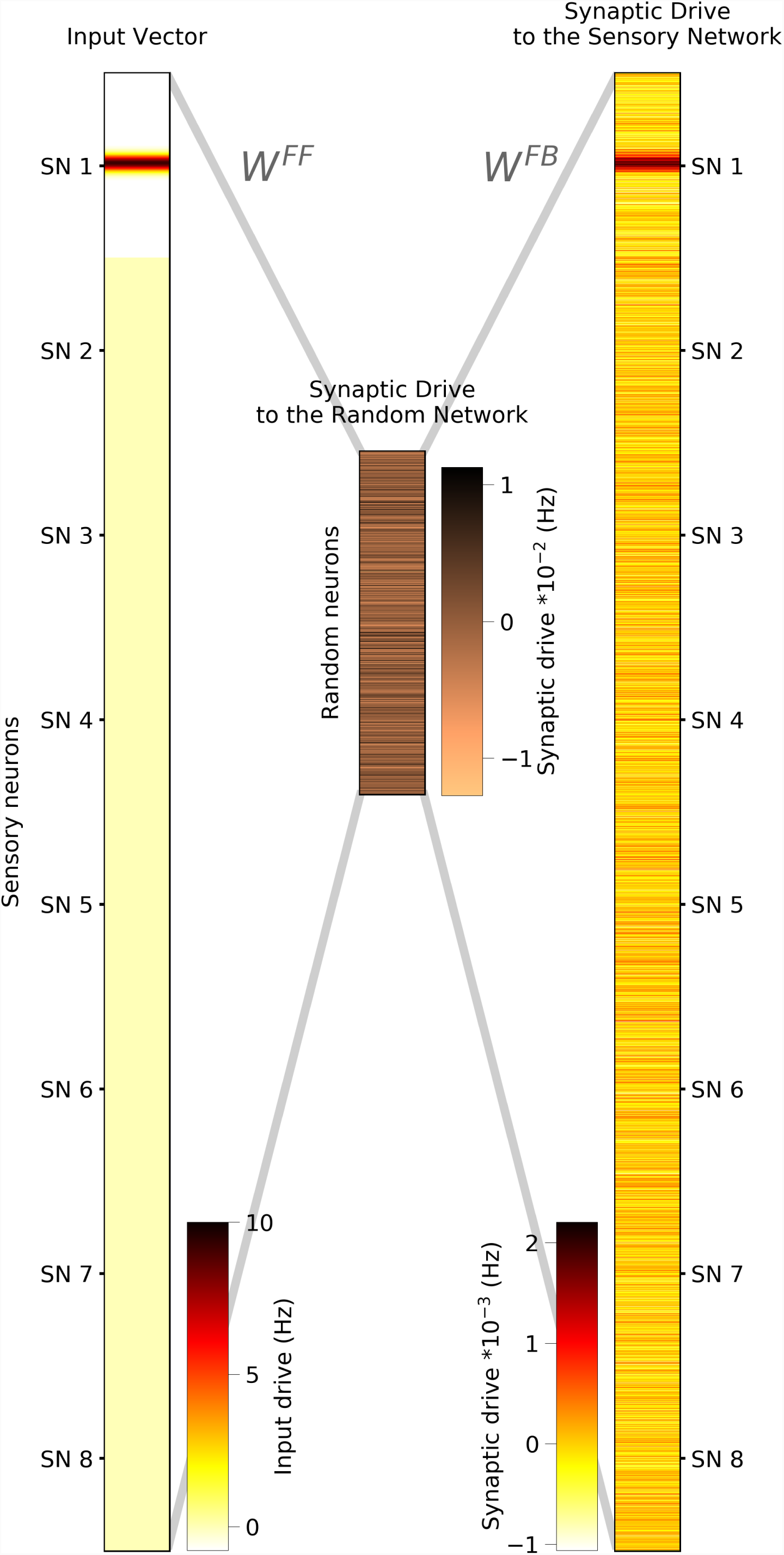
The synaptic drive fed back into a sensory sub-network from the random network closely matches its own representation. Related to Figure 1. **(Left)** Synaptic drive for the entire sensory network resulting from a sensory input into sensory sub-network 1. **(Middle)** Synaptic drive to the random network resulting from the sensory input (left). Estimated by passing synaptic drive from sensory network through *W*^*F*^ ^*F*^. **(Right)** Feedback synaptic drive to the sensory network from the random network. Estimated by passing synaptic drive to random network (middle) through *W*^*FB*^.

**Figure S2.**
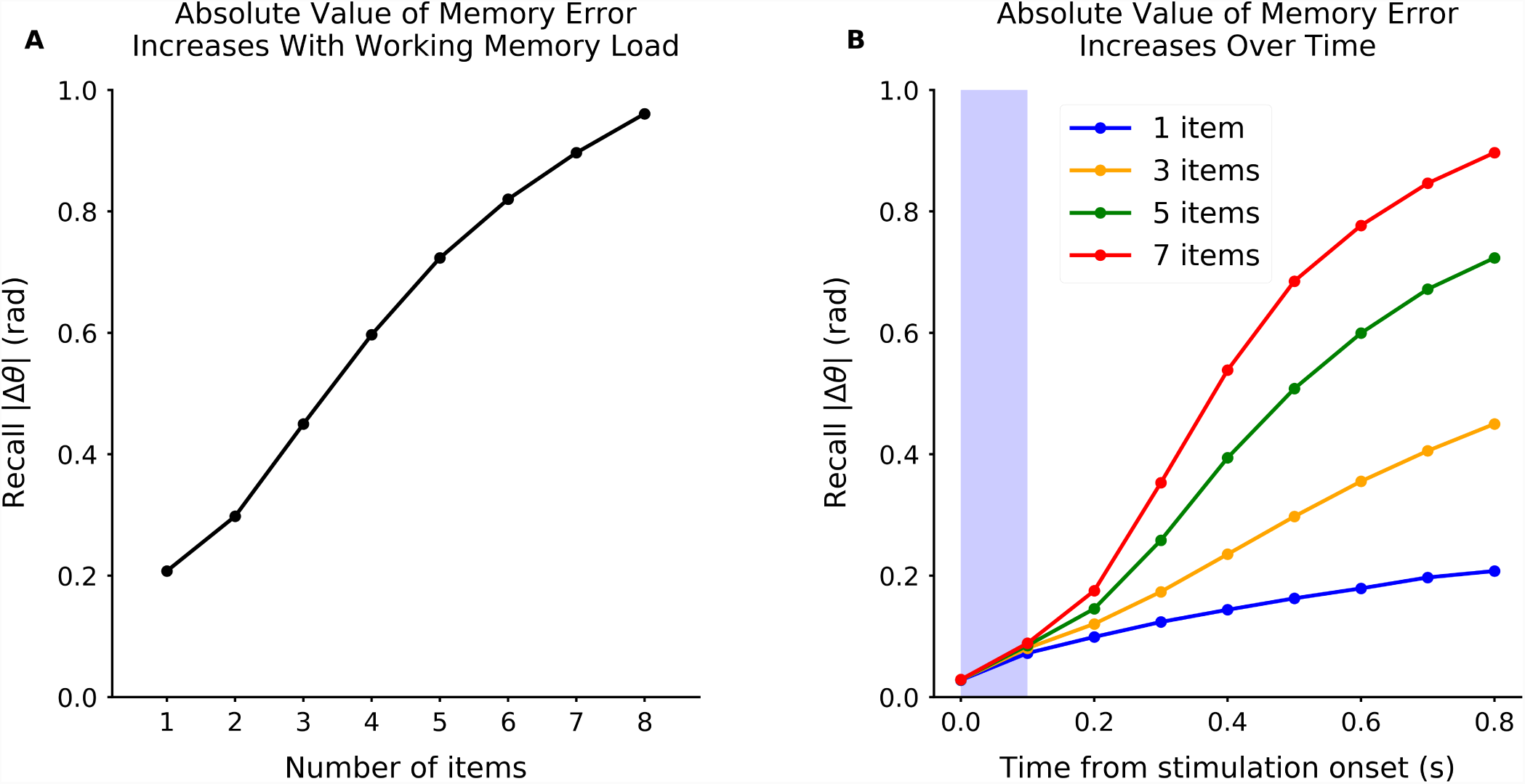
Memory error increases with memory load. Related to Figure 3C,D. Absolute value of the circular error computed from ML spike decoding after 1 second of network simulation, as a function of load **(A)**, and over time **(B)**. Decoding time window is 100msec, forward in time from the indicated time.

**Figure S3.**
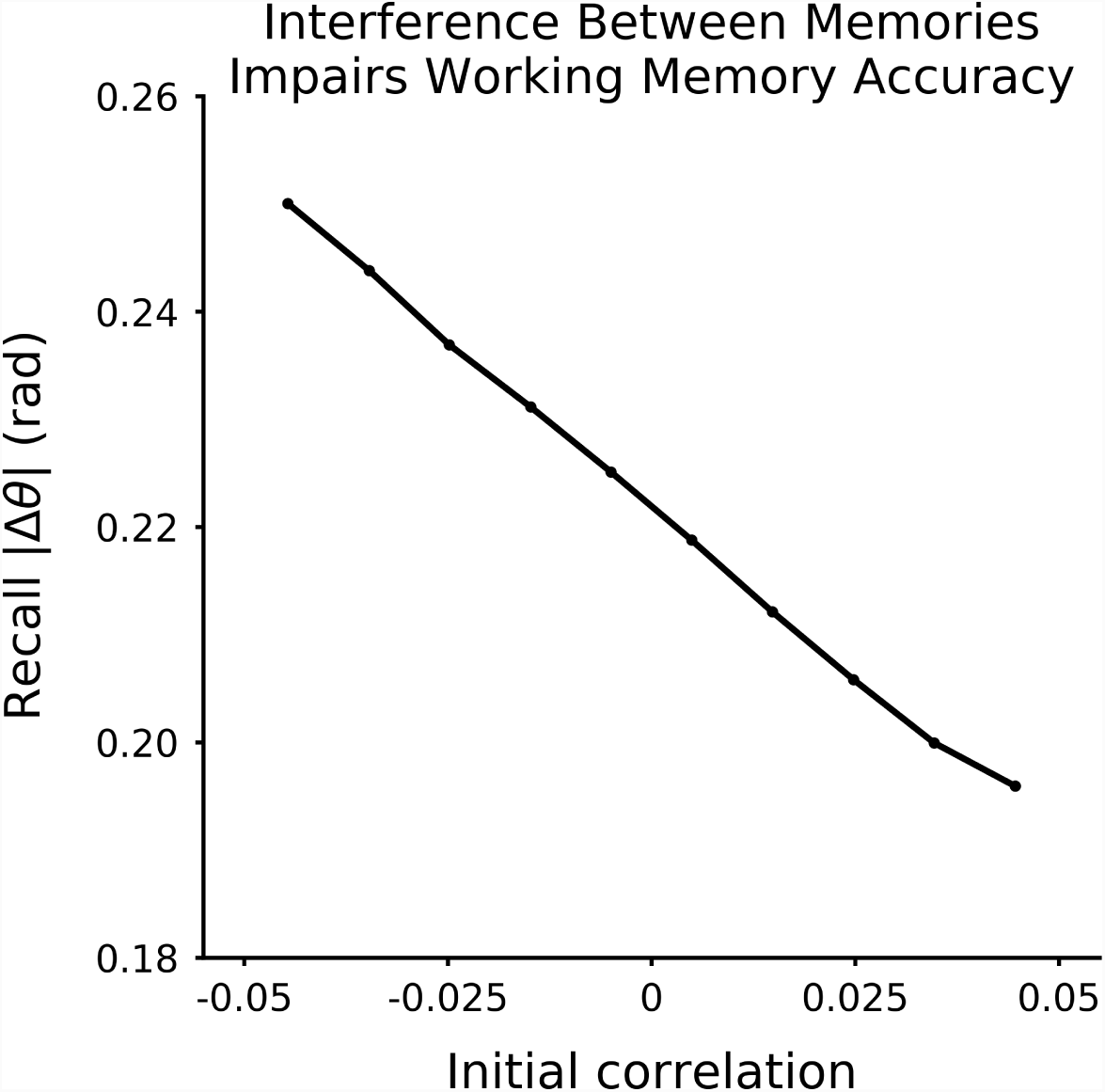
Memory error decreases with reduced interference. Related to Figure 5A. Absolute value of circular error as a function of the correlation between two inputs in two sensory sub-networks. Simulations where both memories are maintained (the error is not due to forgetting).

**Figure S4.**
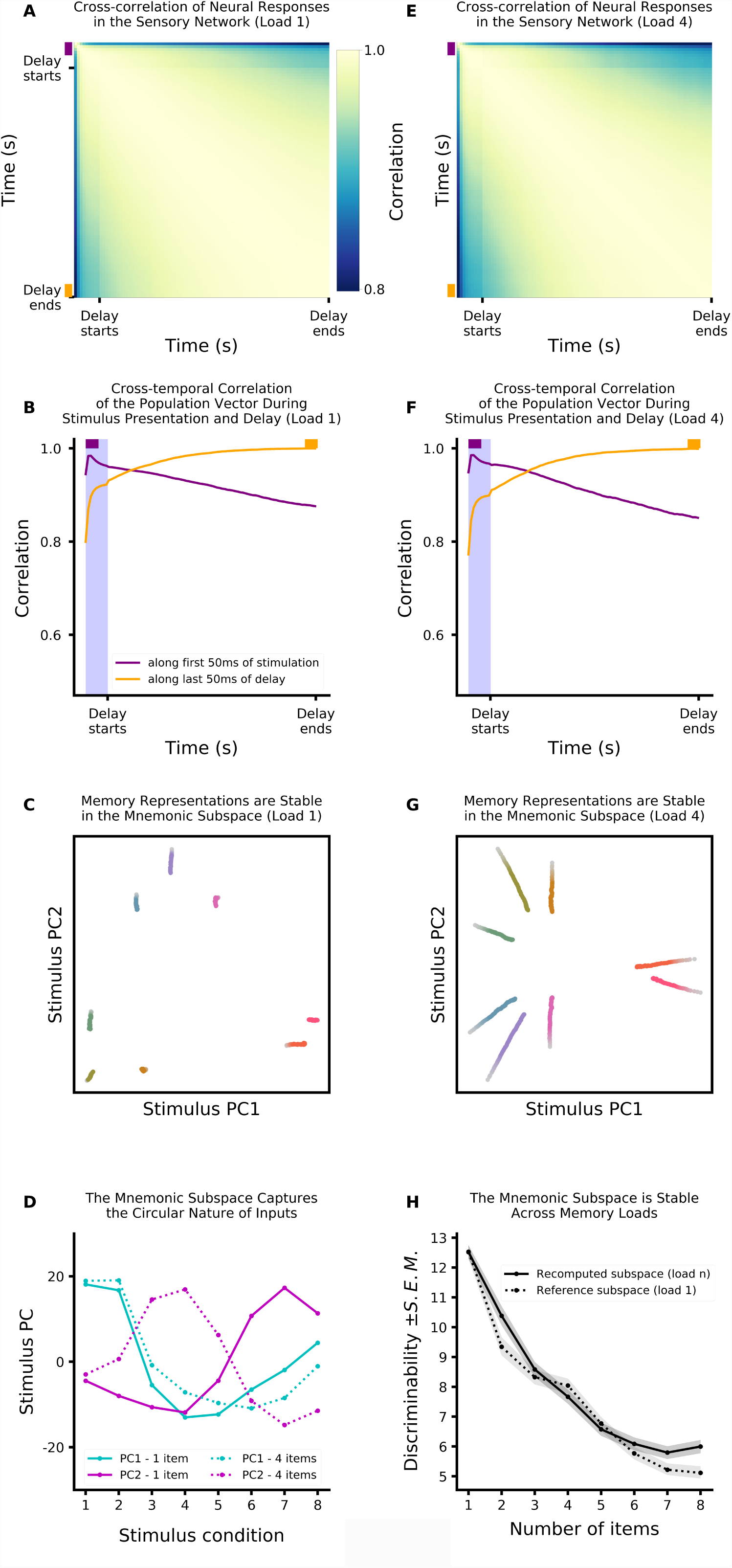
Dynamics in sensory network. PCA analysis of the sensory sub-network activity receiving one of 8 different stimuli (equally separated). As noted in the main text, these simulations used a network with weak direct sensory input into the random network and weak recurrence within the random network (see Methods for details). Figure layout follows Figure 6 (which shows the same for the random network). textbf(A) Cross-correlation of neural activity in the sensory network. Here time-lagged autocorrelation matrix of activity (correlation between the population state at one time point and the state at another time point) shows activity during the stimulation is more correlated with activity during the delay period than for the random network (Fig. 6). **(B)** Slices of the matrix represented in (A): correlation of population state from the first 50ms of the stimulus period (purple) and the last 50ms of the delay period (orange) against all other times. **(C)** The encoding is stable. Here the response of the sensory network population is projected onto the mnemonic subspace (defined by the first two principal components of time-averaged activity, see Methods for details). Each trace corresponds to the response to a different input into sensory sub-network 1, shown over time (from lighter to darker colors). **(D)** Mnemonic subspace is defined by two orthogonal, quasi-sinusoidal representation of inputs, capturing the circular nature of sensory sub-networks. **(E-G)** Same as **A-C** but for a load of 4. Only includes simulations for which the memory in sensory sub-network 1 was maintained (other three memories can be forgotten). **(H)** The mnemonic subspace is stable across working memory load. Decodability of memory (here measured as discriminability, d’, between inputs, see Methods for details) is similar regardless of whether mnemonic subspace was defined for a single input (dashed line) or specific for each load (solid line). Decodability was not significantly different across subspaces for load 3, 4, 5 and 6 (a two-sample Wald test for optimized subspace versus reference subspace gives respectively *p* = 0.45, *p* = 0.28, *p* = 0.56, and *p* = 0.27). For load 2, 7, and 8, decodability was significantly better for the load-specific subspace (*p* = 0.014, *p* = 0.034, and *p* = 0.0019 respectively). Decodability is reduced with load (*p <* 0.001).

**Figure S5.**
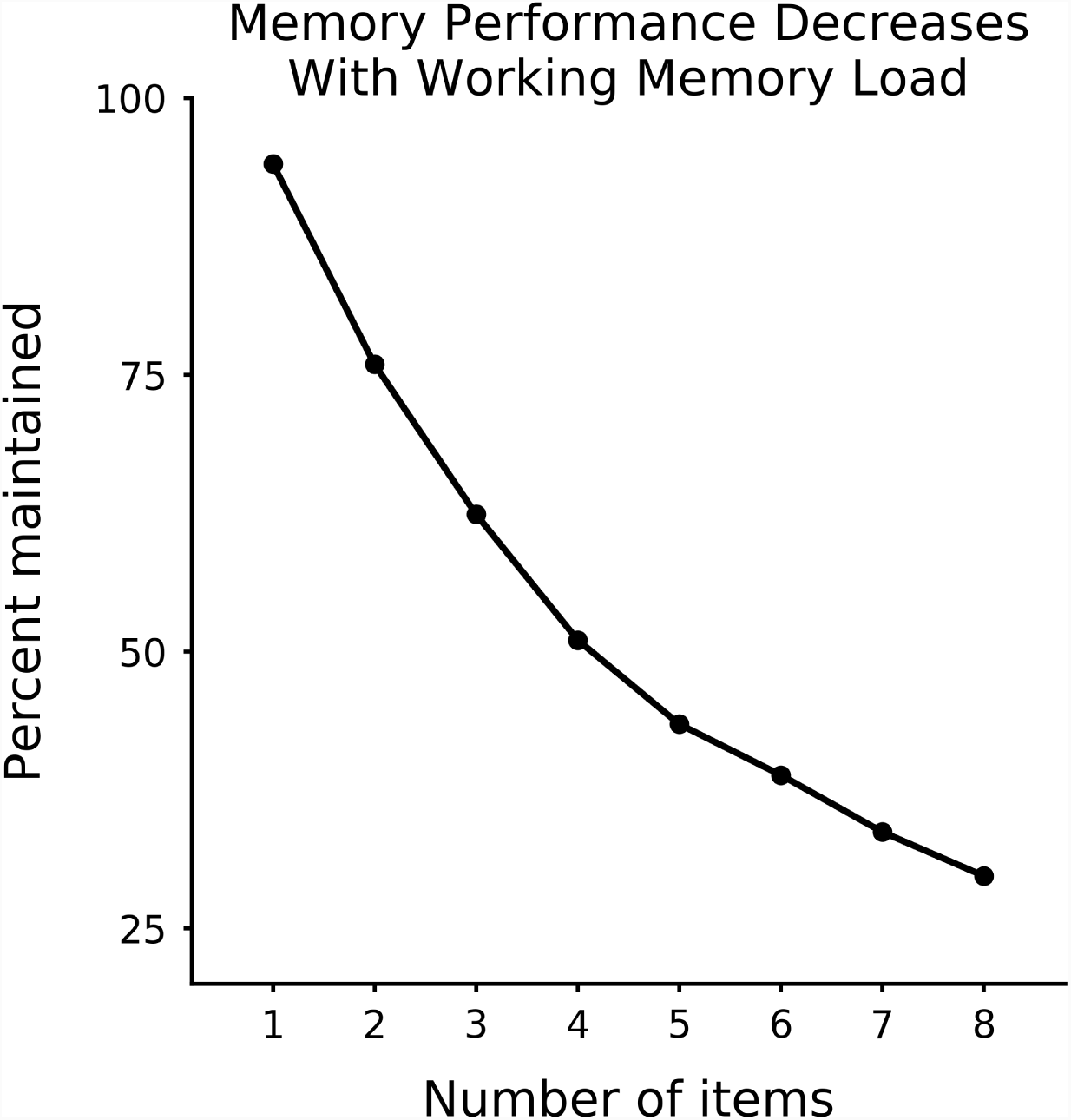
Adding direct sensory inputs into the random network and recurrence within the random network did not alter network performance. Related to Figure 6. Percentage of correct memories, after 1 second of network simulation, as a function of load (as in Fig. 3A). Network included direct transient input into the random network and weak recurrence within the random network (as in Fig. 6).

**Figure S6.**
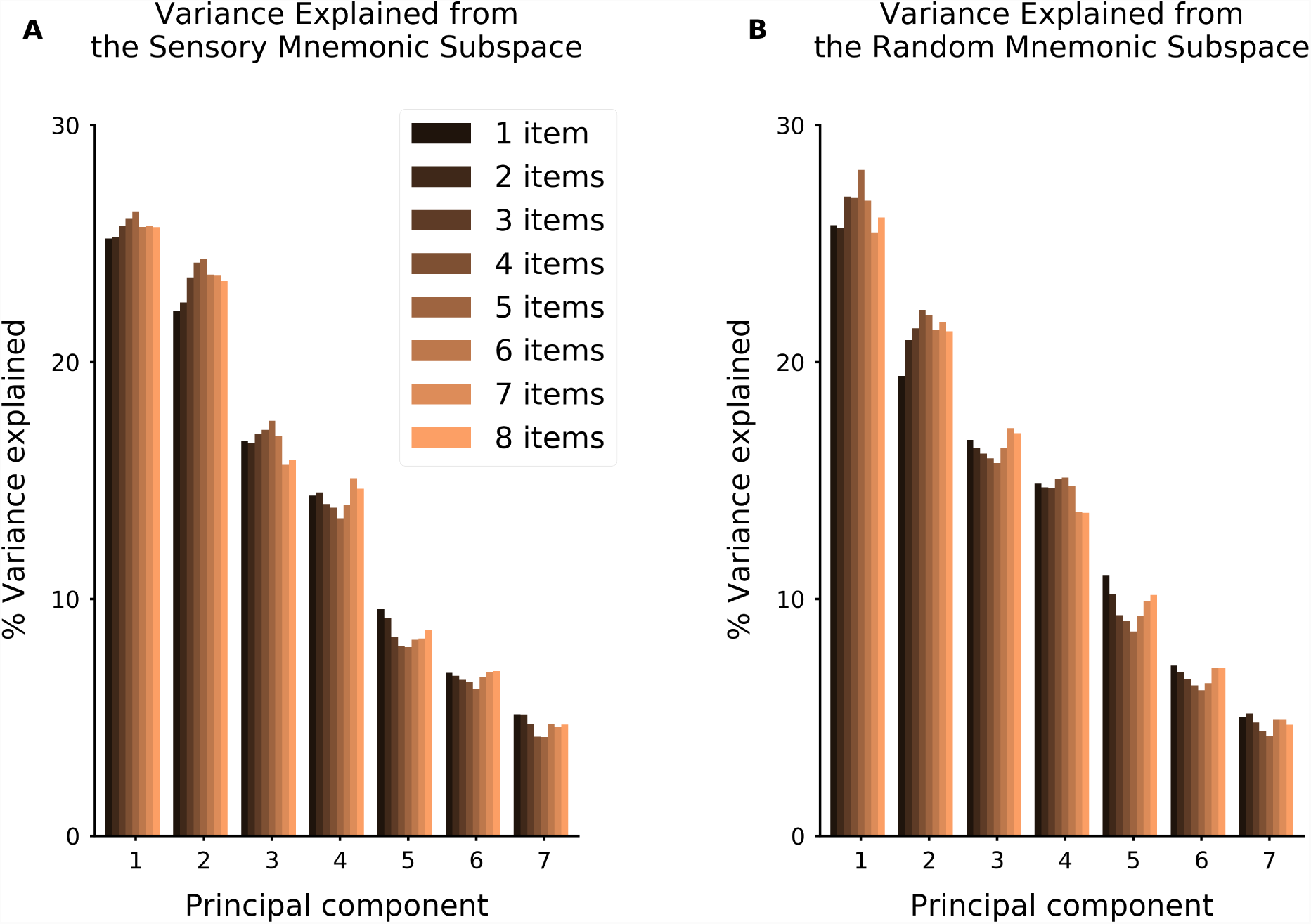
Dimensionality of encoding of a single stimulus is similar across memory load. Related to Figure 6. Percentage of variance explained by each principal axis, for time-averaged delay activity. Computed for sensory sub-network 1 **(A)** and for the random network **(B)**. Follows [Murray et al., 2016].

**Figure S7.**
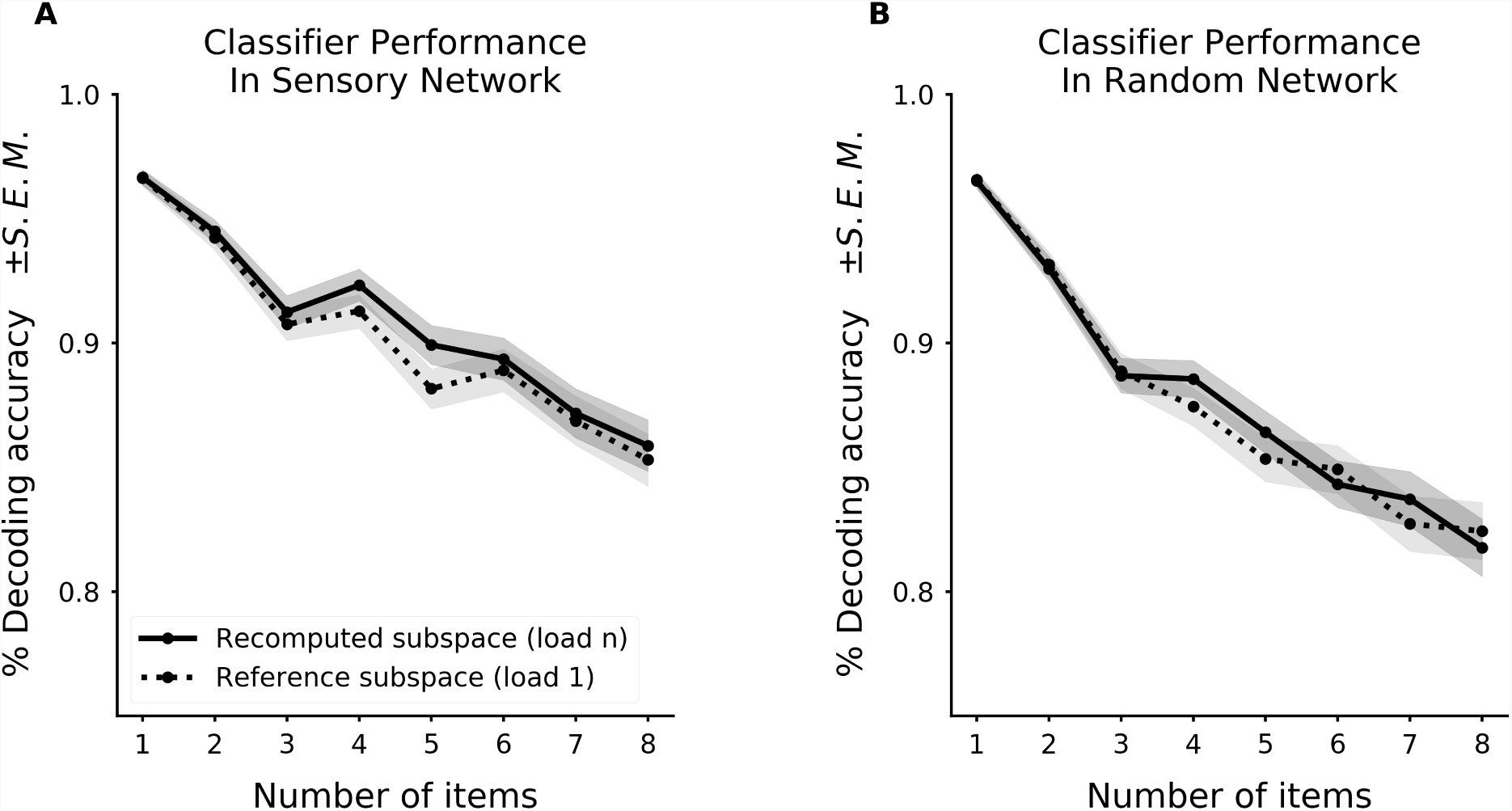
Performance of centroid classifier is similar in the mnemonic subspace defined for any load. Related to Figure 6. Performance of the nearest centroid classifier when the coding subspace was defined for the current load (solid lines) or when the coding subspace was defined with a single item (load 1). Shown for both sensory network (**A**) and random network (**B**). Decodability was not significantly different across subspaces (a two-sample Wald test for load-specific subspace versus reference subspace gives *p >* 0.1 for all loads).

**Figure S8.**
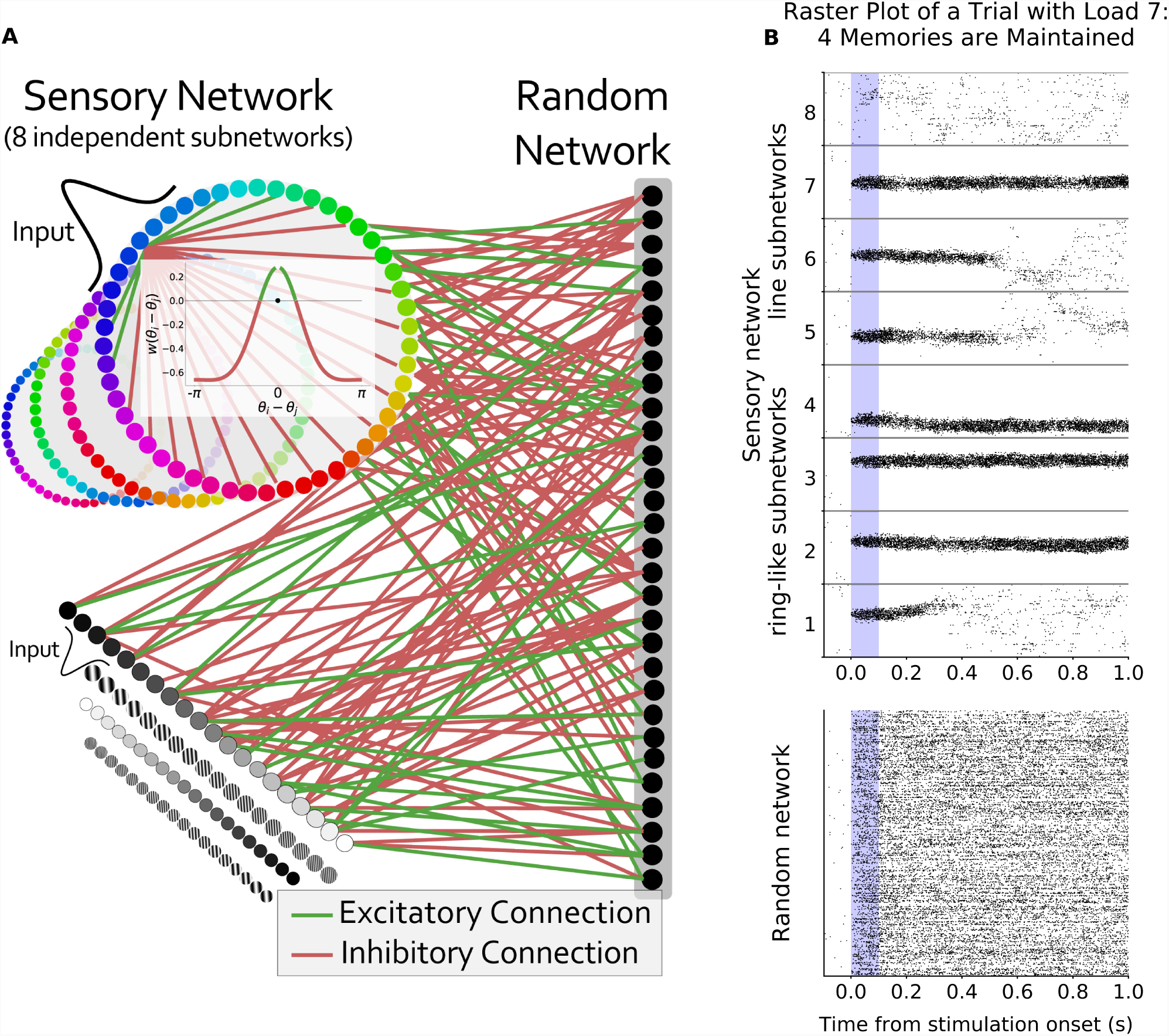
Network performance did not depend on architecture of sensory sub-networks. Related to Figure 1. **(A)** Schematic of alternative model architecture. The sensory network is now composed of 4 ring sub-networks, and 4 line sub-networks. All other parameters remained the same. **(B)** Raster plot of an example trial with 8 sensory sub-networks (512 neurons each) randomly connected to the same random network (1024 neurons). 7 sensory sub-networks (4 ring, 3 line sub-networks) receive a Gaussian input for 0.1 seconds during the ‘stimulus presentation’ period (shaded blue region). As for the original architecture, memories are maintained but the network has a capacity limit.

**Figure S9.**
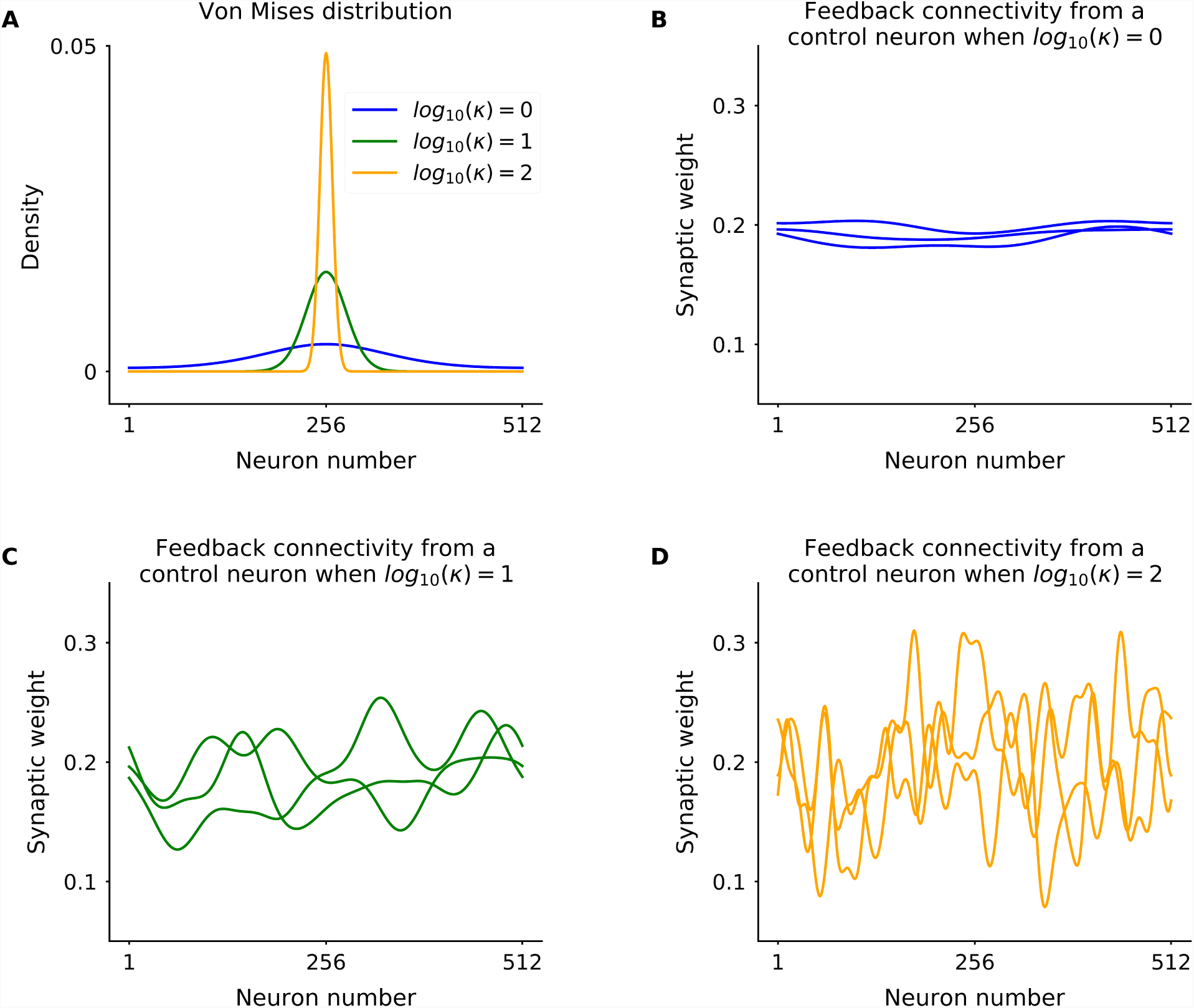
Example Von Mises distributions for spreading of feedback connections. Related to Figure 7. In Fig. 7D, each feedback excitatory connections was distributed according to a Von Mises distribution with varying concentrations. **(A)** Example Von Mises distributions for different concentration parameters. **(B-D)** An example of feedback connectivity as a sum of VonMises from a neuron from the random network to a sensory sub-network (neuron 1 to 512) when **(B)** *κ* = 1, **(C)** *κ* = 10 and **(D)** *κ* = 100.

## References

Alexander, G. (1987). Selective neuronal discharge in monkey putamen reflects intended direction of planned limb movements. Experimental Brain Research, 67(3):623–634.

Atallah, B. V., Bruns, W., Carandini, M., and Scanziani, M. (2012). Parvalbumin-expressing interneurons linearly transform cortical responses to visual stimuli. Neuron, 73(1):159–170.

Barak, O. and Tsodyks, M. (2014). Working models of working memory. Current opinion in neurobiology, 25:20–24.

Barak, O., Tsodyks, M., and Romo, R. (2010). Neuronal population coding of parametric working memory. Journal of Neuroscience, 30(28):9424–9430.

Bays, P. M. (2014). Noise in neural populations accounts for errors in working memory. Journal of Neuroscience, 34(10):3632–3645.

Bays, P. M. (2015). Spikes not slots: noise in neural populations limits working memory. Trends in cognitive sciences, 19(8):431–438.

Bays, P. M., Catalao, R. F., and Husain, M. (2009). The precision of visual working memory is set by allocation of a shared resource. Journal of vision, 9(10):7–7.

Bays, P. M. and Husain, M. (2008). Dynamic shifts of limited working memory resources in human vision. Science, 321(5890):851–854.

Brody, C. D., Romo, R., and Kepecs, A. (2003). Basic mechanisms for graded persistent activity: discrete attractors, continuous attractors, and dynamic representations. Current opinion in neurobiology, 13(2):204–211.

Brunel, N. (2003). Dynamics and plasticity of stimulus-selective persistent activity in cortical network models. Cerebral Cortex, 13(11):1151–1161.

Burak, Y. and Fiete, I. R. (2012). Fundamental limits on persistent activity in networks of noisy neurons. Proceedings of the National Academy of Sciences, 109(43):17645–17650.

Buschman, T. J., Siegel, M., Roy, J. E., and Miller, E. K. (2011). Neural substrates of cognitive capacity limitations. Proceedings of the National Academy of Sciences, 108(27):11252–11255.

Carandini, M. and Heeger, D. J. (1994). Summation and division by neurons in primate visual cortex. Science, 264(5163):1333–1336.

Carandini, M. and Heeger, D. J. (2012). Normalization as a canonical neural computation. Nature Reviews Neuroscience, 13(1):51–62.

Christophel, T. B., Cichy, R. M., Hebart, M. N., and Haynes, J.-D. (2015). Parietal and early visual cortices encode working memory content across mental transformations. Neuroimage, 106:198–206.

Christophel, T. B., Klink, P. C., Spitzer, B., Roelfsema, P. R., and Haynes, J.-D. (2017). The distributed nature of working memory. Trends in Cognitive Sciences, 21(2):111–124.

Compte, A., Brunel, N., Goldman-Rakic, P. S., and Wang, X.-J. (2000). Synaptic mechanisms and network dynamics underlying spatial working memory in a cortical network model. Cerebral Cortex, 10(9):910–923.

Constantinidis, C. and Klingberg, T. (2016). The neuroscience of working memory capacity and training. Nature Reviews Neuroscience, 17(7):438–449.

Constantinidis, C. and Steinmetz, M. A. (1996). Neuronal activity in posterior parietal area 7a during the delay periods of a spatial memory task. Journal of Neurophysiology, 76(2):1352–1355.

Cowan, N. (2010). The magical mystery four: How is working memory capacity limited, and why? Current directions in psychological science, 19(1):51–57.

Curtis, C. E. and D’Esposito, M. (2003). Persistent activity in the prefrontal cortex during working memory. Trends in cognitive sciences, 7(9):415–423.

De Wit, J. and Ghosh, A. (2016). Specification of synaptic connectivity by cell surface interactions. Nature Reviews Neuroscience, 17(1):4.

Druckmann, S. and Chklovskii, D. B. (2012). Neuronal circuits underlying persistent representations despite time varying activity. Current Biology, 22(22):2095–2103.

Duncan, J. (2001). An adaptive coding model of neural function in prefrontal cortex. Nature Reviews Neuroscience, 2(11).

Edin, F., Klingberg, T., Johansson, P., McNab, F., Tegnér, J., and Compte, A. (2009). Mechanism for top-down control of working memory capacity. Proceedings of the National Academy of Sciences, 106(16):6802–6807.

Egorov, A. V., Hamam, B. N., Fransén, E., Hasselmo, M. E., and Alonso, A. A. (2002). Graded persistent activity in entorhinal cortex neurons. Nature, 420(6912):173–178.

Ester, E. F., Sprague, T. C., and Serences, J. T. (2015). Parietal and frontal cortex encode stimulus-specific mnemonic representations during visual working memory. Neuron, 87(4):893–905.

Fransén, E., Tahvildari, B., Egorov, A. V., Hasselmo, M. E., and Alonso, A. A. (2006). Mechanism of graded persistent cellular activity of entorhinal cortex layer v neurons. Neuron, 49(5):735–746.

Funahashi, S., Bruce, C. J., and Goldman-Rakic, P. S. (1989). Mnemonic coding of visual space in the monkey9s dorsolateral prefrontal cortex. Journal of neurophysiology, 61(2):331–349.

Fusi, S., Miller, E. K., and Rigotti, M. (2016). Why neurons mix: high dimensionality for higher cognition. Current opinion in neurobiology, 37:66–74.

Fuster, J. M. (1973). Unit activity in prefrontal cortex during delayed-response performance: neuronal correlates of transient memory. Journal of Neurophysiology.

Fuster, J. M. (1999). Memory in the cerebral cortex: An empirical approach to neural networks in the human and nonhuman primate. MIT press.

Goldman-Rakic, P. S. (1995). Cellular basis of working memory. Neuron, 14(3):477–485.

Goodman, D. F., Stimberg, M., Yger, P., and Brette, R. (2014). Brian 2: eural simulations on a variety of computational hardware. BMC Neuroscience, 15(1):P199.

Harrison, S. A. and Tong, F. (2009). Decoding reveals the contents of visual working memory in early visual areas. Nature, 458(7238):632.

Hasselmo, M. E. and Stern, C. E. (2006). Mechanisms underlying working memory for novel information. Trends in cognitive sciences, 10(11):487–493.

Hebb, D. O. (1949). The organization of behavior: A neuropsychological approach. John Wiley & Sons.

Heeger, D. J. (1992). Normalization of cell responses in cat striate cortex. Visual neuroscience, 9(2):181– 197.

Heeger, D. J., Simoncelli, E. P., and Movshon, J. A. (1996). Computational models of cortical visual processing. Proceedings of the National Academy of Sciences, 93(2):623–627.

Holmgren, C., Harkany, T., Svennenfors, B., and Zilberter, Y. (2003). Pyramidal cell communication within local networks in layer 2/3 of rat neocortex. The Journal of physiology, 551(1):139–153.

Jacobsen, C. F., Elder, J. H., and Haslerud, G. M. (1936). Studies of cerebral function in primates.

Jaeger, H. (2002). Short term memory in echo state networks. gmd-report 152. In GMD-German National Research Institute for Computer Science (2002), http://www.faculty.jacobs-university.de/hjaeger/pubs/STMEchoStatesTechRep.pdf. Citeseer.

Jun, J. K., Miller, P., Hernández, A., Zainos, A., Lemus, L., Brody, C. D., and Romo, R. (2010). Heterogenous population coding of a short-term memory and decision task. Journal of Neuroscience, 30(3):916–929.

Kim, S. S., Rouault, H., Druckmann, S., and Jayaraman, V. (2017). Ring attractor dynamics in the drosophila central brain. Science, 356(6340):849–853.

Kiyonaga, A. and Egner, T. (2016). Center-surround inhibition in working memory. Current Biology, 26(1):64–68.

Ko, H., Hofer, S. B., Pichler, B., Buchanan, K. A., Sjöström, P. J., and Mrsic-Flogel, T. D. (2011). Functional specificity of local synaptic connections in neocortical networks. Nature, 473(7345):87.

Kostadinov, D. and Sanes, J. R. (2015). Protocadherin-dependent dendritic self-avoidance regulates neural connectivity and circuit function. Elife, 4:e08964.

Kuffler, S. W. (1953). Discharge patterns and functional organization of mammalian retina. Journal of neurophysiology, 16(1):37–68.

Levy, R. and Goldman-Rakic, P. S. (1999). Association of storage and processing functions in the dorsolateral prefrontal cortex of the nonhuman primate. Journal of Neuroscience, 19(12):5149–5158.

Loewenstein, Y. and Sompolinsky, H. (2003). Temporal integration by calcium dynamics in a model neuron. Nature neuroscience, 6(9):961–967.

Luck, S. J. and Vogel, E. K. (1997). The capacity of visual working memory for features and conjunctions. Nature, 390(6657):279–281.

Lundqvist, M., Rose, J., Herman, P., Brincat, S. L., Buschman, T. J., and Miller, E. K. (2016). Gamma and beta bursts underlie working memory. Neuron, 90(1):152–164.

Ma, W. J., Husain, M., and Bays, P. M. (2014). Changing concepts of working memory. Nature neuroscience, 17(3):347–356.

Machens, C. K., Romo, R., and Brody, C. D. (2005). Flexible control of mutual inhibition: a neural model of two-interval discrimination. Science, 307(5712):1121–1124.

Mariño, J., Schummers, J., Lyon, D. C., Schwabe, L., Beck, O., Wiesing, P., Obermayer, K., and Sur, M. (2005). Invariant computations in local cortical networks with balanced excitation and inhibition. Nature neuroscience, 8(2):194.

Markram, H., Lübke, J., Frotscher, M., Roth, A., and Sakmann, B. (1997). Physiology and anatomy of synaptic connections between thick tufted pyramidal neurones in the developing rat neocortex. The Journal of physiology, 500(2):409–440.

McKenzie, S., Frank, A. J., Kinsky, N. R., Porter, B., Rivière, P. D., and Eichenbaum, H. (2014). Hippocampal representation of related and opposing memories develop within distinct, hierarchically organized neural schemas. Neuron, 83(1):202–215.

Mendoza-Halliday, D., Torres, S., and Martinez-Trujillo, J. C. (2014). Sharp emergence of feature-selective sustained activity along the dorsal visual pathway. Nature neuroscience, 17(9):1255.

Miller, E. K. and Cohen, J. D. (2001). An integrative theory of prefrontal cortex function. Annual review of neuroscience, 24(1):167–202.

Miller, E. K., Li, L., and Desimone, R. (1993). Activity of neurons in anterior inferior temporal cortex during a short-term memory task. Journal of Neuroscience, 13(4):1460–1478.

Mongillo, G., Barak, O., and Tsodyks, M. (2008). Synaptic theory of working memory. Science, 319(5869):1543–1546.

Murray, J. D., Bernacchia, A., Roy, N. A., Constantinidis, C., Romo, R., and Wang, X.-J. (2016). Stable population coding for working memory coexists with heterogeneous neural dynamics in prefrontal cortex. Proceedings of the National Academy of Sciences, page 201619449.

Packer, A. M. and Yuste, R. (2011). Dense, unspecific connectivity of neocortical parvalbumin-positive interneurons: a canonical microcircuit for inhibition? Journal of Neuroscience, 31(37):13260–13271.

Pasternak, T. and Greenlee, M. W. (2005). Working memory in primate sensory systems. Nature Reviews Neuroscience, 6(2):97.

Pedregosa, F., Varoquaux, G., Gramfort, A., Michel, V., Thirion, B., Grisel, O., Blondel, M., Prettenhofer, P., Weiss, R., Dubourg, V., et al. (2011). Scikit-learn: Machine learning in python. Journal of machine learning research, 12(Oct):2825–2830.

Pertzov, Y., Bays, P. M., Joseph, S., and Husain, M. (2013). Rapid forgetting prevented by retrospective attention cues. Journal of Experimental Psychology: Human Perception and Performance, 39(5):1224.

Pertzov, Y., Manohar, S., and Husain, M. (2017). Rapid forgetting results from competition over time between items in visual working memory. Journal of Experimental Psychology: Learning, Memory, and Cognition, 43(4):528.

Petrides, M. (1995). Impairments on nonspatial self-ordered and externally ordered working memory tasks after lesions of the mid-dorsal part of the lateral frontal cortex in the monkey. Journal of Neuroscience, 15(1):359–375.

Postle, B. R. (2017). Working memory functions of the prefrontal cortex. In The Prefrontal Cortex as an Executive, Emotional, and Social Brain, pages 39–48. Springer.

Pouget, A., Dayan, P., and Zemel, R. (2000). Information processing with population codes. Nature Reviews Neuroscience, 1(2):125.

Rademaker, R. L., Park, Y. E., Sack, A. T., and Tong, F. (2018). Evidence of gradual loss of precision for simple features and complex objects in visual working memory. Journal of Experimental Psychology: Human Perception and Performance.

Rainer, G., Asaad, W. F., and Miller, E. K. (1998). Selective representation of relevant information by neurons in the primate prefrontal cortex. Nature, 393(6685):577.

Rainer, G. and Miller, E. K. (2002). Timecourse of object-related neural activity in the primate prefrontal cortex during a short-term memory task. European Journal of Neuroscience, 15(7):1244–1254.

Rajan, K., Harvey, C. D., and Tank, D. W. (2016). Recurrent network models of sequence generation and memory. Neuron, 90(1):128–142.

Reynolds, J. H., Chelazzi, L., and Desimone, R. (1999). Competitive mechanisms subserve attention in macaque areas v2 and v4. Journal of Neuroscience, 19(5):1736–1753.

Rigotti, M., Barak, O., Warden, M. R., Wang, X.-J., Daw, N. D., Miller, E. K., and Fusi, S. (2013). The importance of mixed selectivity in complex cognitive tasks. Nature, 497(7451):585–590.

Riley, M. R. and Constantinidis, C. (2016). Role of prefrontal persistent activity in working memory. Frontiers in systems neuroscience, 9:181.

Romo, R., Hernández, A., Zainos, A., Lemus, L., and Brody, C. D. (2002). Neuronal correlates of decision-making in secondary somatosensory cortex. Nature neuroscience, 5(11):1217–1225.

Seelig, J. D. and Jayaraman, V. (2015). Neural dynamics for landmark orientation and angular path integration. Nature, 521(7551):186.

Serences, J. T., Ester, E. F., Vogel, E. K., and Awh, E. (2009). Stimulus-specific delay activity in human primary visual cortex. Psychological science, 20(2):207–214.

Seung, H. S., Lee, D. D., Reis, B. Y., and Tank, D. W. (2000). Stability of the memory of eye position in a recurrent network of conductance-based model neurons. Neuron, 26(1):259–271.

Song, S., Sjöström, P. J., Reigl, M., Nelson, S., and Chklovskii, D. B. (2005). Highly nonrandom features of synaptic connectivity in local cortical circuits. PLoS biology, 3(3):e68.

Sprague, T. C., Ester, E. F., and Serences, J. T. (2014). Reconstructions of information in visual spatial working memory degrade with memory load. Current Biology, 24(18):2174–2180.

Stimberg, M., Goodman, D. F., Benichoux, V., and Brette, R. (2013). Brian 2-the second coming: spiking neural network simulation in python with code generation. BMC Neuroscience, 14(1):P38.

Stokes, M. G. (2015). ‘activity-silent’ working memory in prefrontal cortex: a dynamic coding framework. Trends in cognitive sciences, 19(7):394–405.

Stokes, M. G., Buschman, T. J., and Miller, E. K. (2017). Cognitive control. The Wiley Handbook of Cognitive Control, page 221.

Stokes, M. G., Kusunoki, M., Sigala, N., Nili, H., Gaffan, D., and Duncan, J. (2013). Dynamic coding for cognitive control in prefrontal cortex. Neuron, 78(2):364–375.

Swan, G. and Wyble, B. (2014). The binding pool: A model of shared neural resources for distinct items in visual working memory. Attention, Perception, & Psychophysics, 76(7):2136–2157.

Vogel, E. K. and Machizawa, M. G. (2004). Neural activity predicts individual differences in visual working memory capacity. Nature, 428(6984):748.

Vogels, T. P. and Abbott, L. F. (2005). Signal propagation and logic gating in networks of integrate-and-fire neurons. Journal of neuroscience, 25(46):10786–10795.

Vogels, T. P., Rajan, K., and Abbott, L. F. (2005). Neural network dynamics. Annu. Rev. Neurosci., 28:357–376.

Vogels, T. P., Sprekeler, H., Zenke, F., Clopath, C., and Gerstner, W. (2011). Inhibitory plasticity balances excitation and inhibition in sensory pathways and memory networks. Science, 334(6062):1569–1573.

Wang, X.-J. (2001). Synaptic reverberation underlying mnemonic persistent activity. Trends in neurosciences, 24(8):455–463.

Watanabe, Y. and Funahashi, S. (2012). Thalamic mediodorsal nucleus and working memory. Neuroscience & Biobehavioral Reviews, 36(1):134–142.

Wilken, P. and Ma, W. J. (2004). A detection theory account of change detection. Journal of vision, 4(12):11–11.

Wilson, N. R., Runyan, C. A., Wang, F. L., and Sur, M. (2012). Division and subtraction by distinct cortical inhibitory networks in vivo. Nature, 488(7411):343.

Yamagishi, T., Yoshitake, K., Kamatani, D., Watanabe, K., Tsukano, H., Hishida, R., Takahashi, K., Takahashi, S., Horii, A., Yagi, T., et al. (2018). Molecular diversity of clustered protocadherin-*a* required for sensory integration and short-term memory in mice. Scientific reports, 8(1):9616.

Zaksas, D. and Pasternak, T. (2006). Directional signals in the prefrontal cortex and in area mt during a working memory for visual motion task. Journal of Neuroscience, 26(45):11726–11742.

Zhang, W. and Luck, S. J. (2008). Discrete fixed-resolution representations in visual working memory. Nature, 453(7192):233–235.

